# Absence of c-Maf and IL-10 enables Type I IFN enhancement of innate responses to low-dose LPS in alveolar macrophages

**DOI:** 10.1101/2024.05.22.594428

**Authors:** Pamelia N. Lim, Maritza M. Cervantes, Linh K. Pham, Sydney Doherty, Ankita Tufts, Divya Dubey, Dat Mai, Alan Aderem, Alan H. Diercks, Alissa C. Rothchild

## Abstract

Alveolar macrophages (AMs) are lower-airway resident myeloid cells and are among the first to respond to inhaled pathogens. Here, we interrogate AM innate sensing to Pathogen Associated Molecular Patterns (PAMPs) and determine AMs have decreased responses to low-dose LPS compared to other macrophages, as measured by TNF, IL-6, *Ifnb*, and *Ifit3*. We find the reduced response to low-dose LPS correlates with minimal TLR4 and CD14 surface expression, despite sufficient internal expression of TLR4. Additionally, we find that AMs do not produce IL-10 in response to a variety of PAMPs due to low expression of transcription factor c-Maf and that lack of IL-10 production contributes to an enhancement of pro-inflammatory responses by Type I IFN. Our findings demonstrate that AMs have cell-intrinsic dampened responses to LPS, which is enhanced by type I IFN exposure. These data implicate conditions where AMs may have reduced or enhanced sentinel responses to bacterial infections.

**HIGHLIGHTS:**

- Alveolar macrophages (AMs) do not produce TNF or IL-6 in response to low-dose LPS due to minimal surface expression of TLR4 and CD14
- Lack of AM IL-10 production is dependent on low c-Maf expression
- Exogenous c-Maf expression increases AM IL-10 production
- IFNβ enhances AM TNF and IL-6 responses to low-dose LPS and this is dependent on a lack of IL-10

## INTRODUCTION

Alveolar macrophages (AMs) are located within lung alveoli and serve as sentinels for inhaled pathogens and airborne environmental particles. AM’s constant exposure to foreign material and concurrent role in pulmonary homeostasis creates a unique profile that shapes their innate response. AMs must mount responses to pathogens they encounter as well as maintain airway clearance through the removal of surfactant and debris. In a healthy lung, AMs are the most abundant myeloid cell^1,2^ and are present in only 1 out of every 3 alveoli^3,4^, so their initial responses to pathogens must be generated in an isolated manner.

AMs participate in innate responses to a variety of inhaled pathogens, including viral infections such as SARS-CoV-2^5,6^, respiratory syncytial virus (RSV)^7^, influenza^8–11^, and Newcastle disease virus (NDV)^12^, where they contribute to inflammatory responses and disease pathogenesis. In contrast, AM responses to bacterial infections are more varied. AMs infected *ex vivo* with *Pseudomonas aeruginosa* produce TNF and IL-6^13^, as well as IL-1β following inflammasome activation^14^. Additionally, depletion of AMs *in vivo* prior to *P. aeruginosa* infection reduces neutrophil recruitment^15^. In contrast, AMs infected *in vivo* with *Legionella pneumophilia* do not produce TNF^16^. AMs are the first cells infected in the lung during *Mycobacterium tuberculosis* (Mtb) infection^17,18^. Mtb-infected AMs mount a cell-protective, NRF2-dependent response with minimal expression of pro-inflammatory genes^18^. We sought to address the discrepancy in AM inflammatory responses by directly examining AM cell-intrinsic innate sensing pathways and how they might be influenced by exogenous signals, such as Type I IFN, which are present in the lung milieu during viral infection.

Innate recognition of lipopolysaccharide (LPS), a critical component of the outer membrane of gram-negative bacteria, is one of the most well-studied sensing pathways. LPS is sensed on the cell surface by TLR4 and CD14, leading to MyD88 and TIRAP signaling, activation of NF-κB, and production of pro-inflammatory cytokines^19–21^. In addition, CD14 binding to TLR4 can result in TLR4 endocytosis, endosomal signaling through TRIF and TRAM adaptors, activation of IRF3, and production of Type I IFN^19,22^. These major pathways have been primarily identified through studies in bone marrow-derived macrophages (BMDMs). An additional study showed the requirement of CD14 for TNF production in response to low-dose LPS in mouse peritoneal macrophages^23^ and peripheral blood mononuclear cells^24^. Yet, it is unknown whether AMs similarly mount a robust response to LPS and other PAMPs under isolated conditions. A better understanding of AM direct innate sensing capacity is critical for studying the role of AMs as airway sentinels during pulmonary infections.

To examine direct, individual AM innate responses to LPS, we delivered LPS linked to microspheres, which could be tracked by fluorescence either *in vivo* by aerosol or *ex vivo* to cultured AMs, using *in vitro* BMDMs as a positive control. We observed that both *in vivo* and *ex vivo* AMs have a significantly reduced inflammatory response to LPS-coated beads compared to BMDMs. Similarly, AMs produce significantly less TNF and IL-6 in response to low-dose soluble LPS, compared to that of BMDMs. This response is PAMP-specific, as AMs mount robust pro- inflammatory and Type I IFN responses to other PAMPs, including Pam3Cys, R848, and 2’3’- cGAMP. We find that while AMs express *Tlr4* mRNA, AMs have minimal surface expression of TLR4 and this is associated with low levels of expression of co-receptor CD14. Interestingly, AMs do not produce IL-10 in response to any PAMPs tested. We show that the lack of AM IL-10 production is dependent on the absence of the transcription factor c-Maf. AM IL-10 production can be rescued by exogenous expression of c-Maf. Lastly, we demonstrate that addition of recombinant IFNβ enhances AM production of TNF and IL-6 in response to LPS, and this is again dependent on the absence of IL-10 production. Overall, our results demonstrate that AMs have PAMP-specific, cell-intrinsic deficiencies in innate sensing, which make them uniquely tolerant to LPS and sensitive to Type I IFN.

## RESULTS

### Alveolar macrophages do not mount a pro-inflammatory response to LPS-conjugated beads *in vivo* and *ex vivo*

To characterize AM responses to LPS and to distinguish direct PAMP sensing from paracrine signaling effects *in vivo*, we coated 1.0 μm polystyrene beads with LPS and delivered them to mice via aerosol. The same preparation of LPS-conjugated beads were delivered in parallel to BMDMs *in vitro* (**Fig. 1A**). This approach was optimized so that 10% or fewer macrophages received LPS-coated beads under each condition, minimizing indirect bystander effects. Both Bead+ (LPS_bead_) and Bead- (bystander) macrophages were sorted from each population (**Fig. S1A, B**) and gene expression was profiled by RNA-sequencing. In agreement with published studies, Gene Set Enrichment Analysis demonstrated that LPS_bead_ BMDMs were significantly enriched for the HALLMARK pathways “Inflammatory response”, “TNFA signaling”, and “IL6 Jak Stat3 signaling” (**Fig. 1B**). Both bystander BMDMs and vehicle control Bead+ BMDMs (PBS_bead_), showed minimal enrichment for these pathways, confirming the specificity of the approach. In contrast, gene expression profiles for LPS_bead_ AMs were not enriched for these pro-inflammatory pathways. Additionally, analysis of Differentially Expressed Genes (DEGs) between LPS_bead_ AM and LPS_bead_ BMDMs demonstrated that only LPS_bead_ BMDMs up-regulated key pro-inflammatory response genes *Tnf* and *Il1b*, and interferon-stimulated genes *Irg1* and *Ch25h* (**Fig. 1C**).

**Figure 1:**
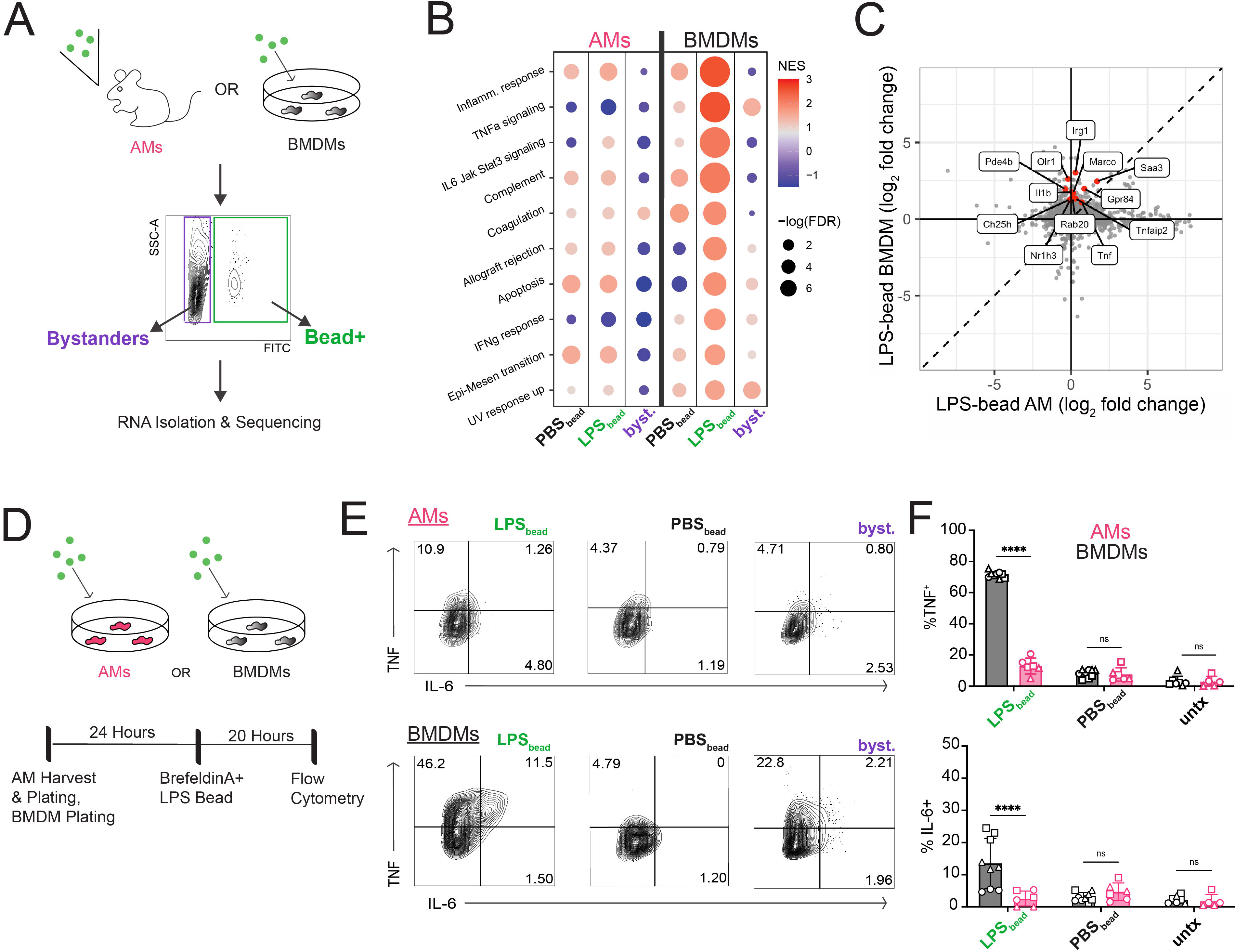
Alveolar macrophages do not mount a pro-inflammatory response to LPS- conjugated beads *in vivo* or *ex vivo*. (A) Schematic representation of LPS-coated beads delivered to mice by nebulization or to BMDMs. Uncoated beads resuspended in PBS (PBS bead) delivered to controls. Cells were isolated and sorted into Bead+ or Bystander groups. (B) Gene Set Enrichment Analysis of the top ten differentially expressed pathways between LPS_Bead_ BMDMs and LPS_Bead_ AMs. (C) Scatter plot of log_2_ fold change values for AM LPS_Bead_ and BMDM LPS_Bead_ populations. Labeled genes are significantly up-regulated (FDR < 0.05, FC > 2) in only the BMDM LPS_Bead_ population. (D) Schematic representation of AMs or BMDMs stimulated with LPS-coated beads *ex vivo* and subsequently analyzed by flow cytometry for intracellular TNF and IL-6. (E) Representative dot plots of the conditions for BMDMs and AMs. (F) Intracellular Cytokine Staining of TNF and IL-6 for LPS_bead_, PBS_bead_, or untreated BMDMs and AMs. Data are representative of 3 independent experiments (E) or compiled from 2 independent experiments (B, C) or 3 independent experiments (F). Technical replicates within each experiment represented by unique shapes. ****P < 0.0001, ns not significant by two-way ANOVA with Sidak’s multiple comparison test.

To determine whether this hypo-response was due to the influence of the lung environment or due to cell-intrinsic regulation, we repeated this experiment with AMs isolated from the lung via bronchoalveolar lavage (BAL) and cultured *ex vivo* for <24 hours. Following 20 hours of exposure to LPS- or PBS-coated beads, TNF and IL-6 production was measured by intracellular cytokine staining (ICS) in the presence of Brefeldin A (**Fig. 1D, Fig. S1C, D**). Across three independent experiments, 10 ± 3.7% of BMDMs and 6.2 ± 2.1% of AMs (mean ± SEM) were Bead+ (**Fig. S1E**). Again, we observed that AM pro-inflammatory responses were significantly diminished compared to BMDMs. 12.9 ± 5.1% of LPS_bead_ AMs were TNF+ compared to 71.4 ± 2.1% of BMDMs. 2.6 ± 2.3% of LPS_bead_ AMs were IL-6+ compared to 13.5 ± 7.7% of BMDMs (**Fig. 1E, F)**. Parallel experiments using peritoneal macrophages (PMs) led to similar cytokine production as observed for BMDMs (**Fig. S1F**), suggesting that the lack of response in AMs is not a general feature of all tissue-resident macrophages. Overall, these data demonstrate that in a scenario where cells directly sense LPS and there is an absence of paracrine signaling, AMs are unable to mount a pro-inflammatory response either *in vivo* or *ex vivo*. These data also show that removal from the lung environment does not enhance AM sensing of low-dose LPS and suggest that the factors that regulate AM sensing of LPS are cell-intrinsic.

### Alveolar macrophages have a high LPS-specific activation threshold

We next sought to evaluate the AM response to soluble LPS which, unlike the bead-conjugated approach, would allow for paracrine signaling and cross-talk between neighboring cells. For monocyte-derived macrophages, paracrine signaling has been shown to be important for the production of significant amounts of IL-6, TNF, and IL-10 in response to LPS stimulation^25^. We measured TNF and IL-6 protein secretion by AMs and BMDMs following stimulation with 0.1, 1, and 10 ng/ml LPS. There was significantly less production of both TNF and IL-6 in AMs compared to BMDMs after stimulation with 1 ng/mL of LPS, but AMs produced more TNF and comparable IL-6 to BMDMs at 10 ng/mL (**Fig. 2A**). To investigate if decreased TNF and IL-6 production was specific to LPS or shared across other ligands, we stimulated AMs and BMDMs with two other PAMPs, Pam3Cys (TLR1:2 agonist) or R848 (TLR7/8 agonist), across 100-fold dose curves. In contrast to our results with LPS, AMs made significantly more TNF and IL-6 than BMDMs for both Pam3Cys (**Fig. S2A**) and R848 (**Fig. S2B**) for most doses tested. Additionally, to determine if AM’s diminished response was shared across other tissue-resident macrophage populations, we repeated the experiments in **Fig. 2A** with peritoneal macrophages (PMs). In contrast to AMs, PMs produced significant levels of TNF and IL-6 starting at 1 ng/mL of LPS (**Fig. S2C**). Additionally, PMs stimulated with Pam3Cys had similar TNF expression to BMDMs and higher IL-6 expression compared to BMDMs and AMs (**Fig. S2D**).

**Figure 2:**
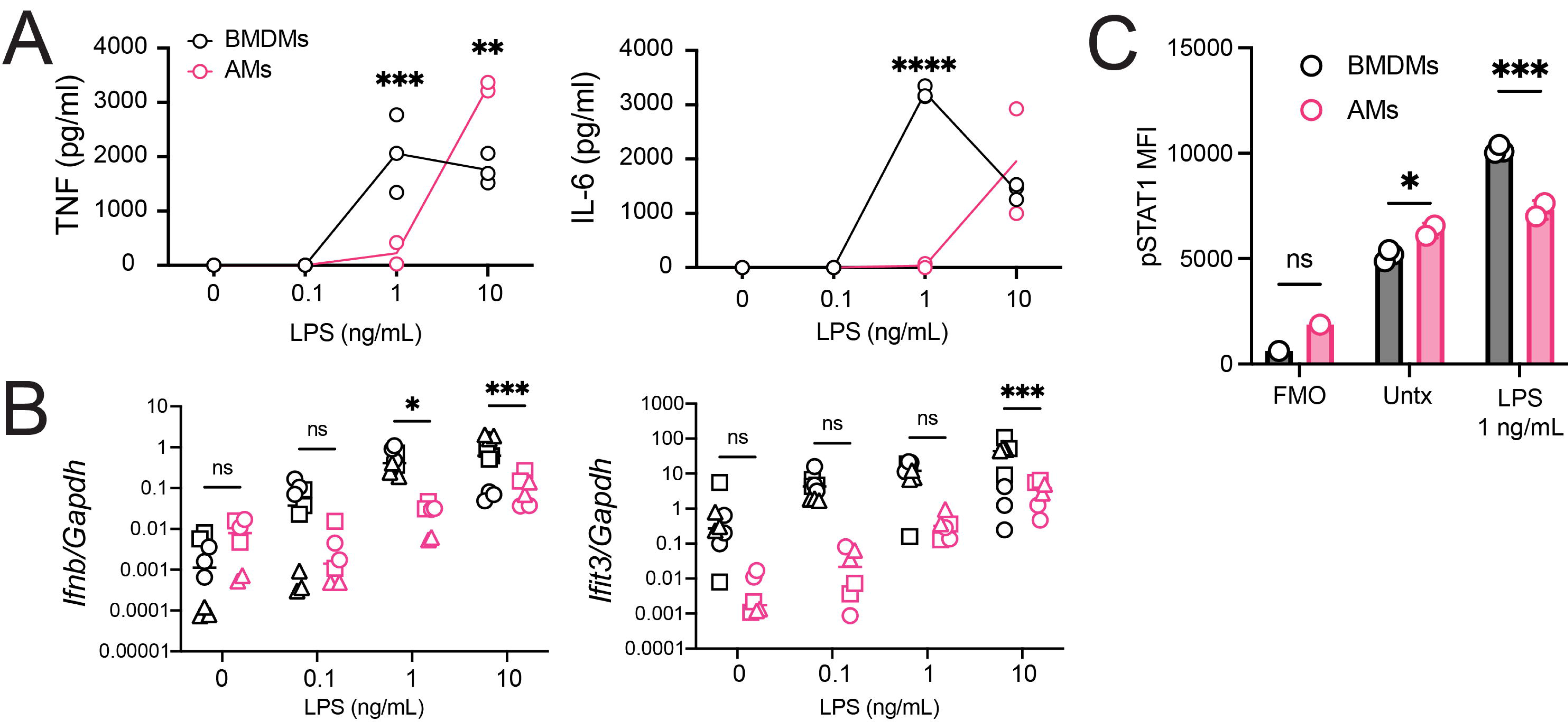
Alveolar macrophages have a high LPS-specific activation threshold. (A) TNF (*left*) and IL-6 (*right*) for AMs (pink, technical duplicate) and BMDMs (black, technical triplicate) either untreated or stimulated with 0.1, 1, or 10 ng/mL of LPS for 20 hours (B) *Ifnb* and *Ifit3* gene expression for 0 – 10 ng/mL LPS for BMDMs (black) and AMs (pink) after 4 hours. (C) phospho- STAT1 intracellular staining MFI of BMDMs (black, technical triplicate) or AMs (pink, technical duplicate) after 20 hours of LPS (1 ng/mL) or fresh media (untreated). Data are representative of 3 independent experiments (A, C) or compiled from 3 independent experiments (B). Technical replicates within each experiment represented by unique shapes. (A-C) *P<0.05, ***P<0.001, ****P<0.0001, ns is not significant, Two-way ANOVA with Sidak’s multiple comparison test.

TLR4 engagement leads to the production of both pro-inflammatory cytokines via MYD88/TIRAP signaling on the cell surface and Type I IFN via TRIF/TRAM signaling within endosomes^20^. To test if LPS signaling through the TRIF/TRAM pathway was also deficient in AMs, we quantified *Ifnb and Ifit3* expression by RT-qPCR across the 100-fold dose curve of LPS. We saw significantly less *Ifnb* expression in AMs after stimulation with 1 ng/mL and 10 ng/mL LPS compared to BMDMs and significantly decreased expression of *Ifit3, an* Interferon Stimulated Gene (ISG), in AMs stimulated with 10 ng/mL LPS (**Fig. 2B**). IFNβ is sensed by the Interferon-α/β receptor (IFNAR) which leads to the activation of STAT1. Measuring phosphorylation of STAT1 by flow cytometry following 1 ng/mL LPS stimulation, we observed that BMDMs had a significant increase in pSTAT1, while AMs showed no increase in pSTAT1 levels over untreated controls (**Fig. 2C**).

To test if decreased AM expression of *Ifnb and Ifit3* was also PAMP-specific, AMs and BMDMs were stimulated with 2’3’-cGAMP, a ligand for STING. Overall, both AMs and BMDMs robustly increased *Ifnb* and *Ifit3* gene expression in response to 2’3’-cGAMP. AMs and BMDMs had similar expression levels of *Ifnb* after stimulation with 0.1 and 1 μM, while BMDMs expressed significantly higher *Ifnb* at the 10 μM dose (**Fig. S2E**). AMs expressed significantly less *Ifit3* after stimulation with 0.1 μM 2’3’-cGAMP compared to BMDMs, but had similar expression levels at higher doses (**Fig. S2E**). These data demonstrate that AMs have a higher LPS-specific activation threshold for TNF, IL-6 and Type I IFN signaling than BMDMs or peritoneal macrophages.

### Low alveolar macrophage TLR4 surface expression is associated with low expression of co-receptor CD14

One potential reason for AM’s higher activation threshold to LPS is reduced expression of either the receptor or signaling adaptors for LPS sensing. We examined AM and BMDM gene expression for TLR4 and associated adaptor proteins. We found increased expression of *Tlr4* and associated adaptors *Myd88*, *Tirap*, and *Ticam2* (Tram) in AMs compared to BMDMs, but significantly lower expression of *Cd14* in AMs compared to BMDMs (**Fig. 3A**). To determine if differences in gene expression were reflected in protein expression, we quantified TLR4 and CD14 surface expression by flow cytometry before and after stimulation with 1 ng/mL LPS. Surprisingly, AMs had significantly lower expression of surface TLR4 at all timepoints, with minimal changes following LPS stimulation (**Fig. 3B**). BMDMs also had a significant decrease in TLR4 surface expression at 4 hours, which aligns with what has been previously reported about the mechanism and timing of TLR4 endosomal recycling^19^. AMs also had significantly lower surface expression of CD14, although both AMs and BMDMs showed increases in surface CD14 at 4 hours post-LPS stimulation (**Fig. 3C**). We also examined TLR4 and CD14 cell surface levels in PMs. PMs showed a similar pattern to BMDMs, with high TLR4 and CD14 expression and a decrease in surface TLR4 expression after LPS stimulation (**Fig. S3C, D**). We additionally measured TLR2 levels under basal conditions and saw similar levels in AMs compared to BMDMs (**Fig. S3E**).

**Figure 3:**
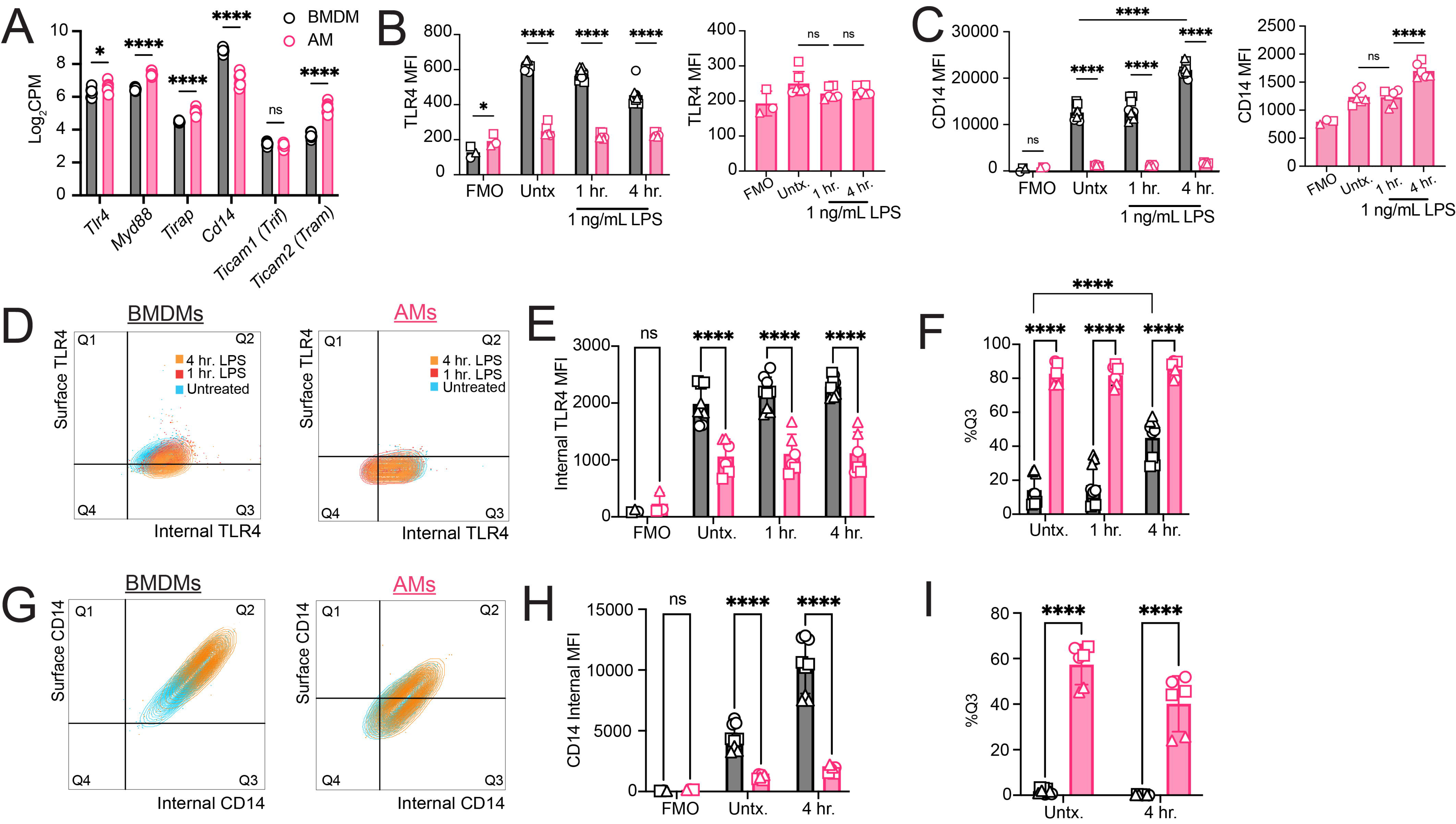
Alveolar macrophages have low surface expression of TLR4 and CD14 compared to other cell types. (A) Gene expression (log_2_ CPM) of AMs (pink) and BMDMs (black) collected in Fig. 1A. (B) TLR4 MFI of AMs (pink, graphed independently to the right) and BMDMs (black) after 0, 1, and 4 hours of LPS (1 ng/mL) stimulation *ex vivo*. (C) CD14 MFI under the same conditions as (B). (D) Surface and internal TLR4 expression of BMDMs and AMs after no treatment (blue), 1 hour (red) or 4 hours (orange) post-LPS stimulation (1 ng/mL). Gates set based on cell-specific FMOs. (E) Internal TLR4 MFI for both BMDMs (black) and AMs (pink). (F) Frequency of Q3^+^ (internal TLR4^+^) BMDMs and AMs. (G) Surface and internal CD14 expression of AMs and BMDMs before and after stimulation with 4 hr. LPS (1 ng/mL). Gates set based on cell-specific FMOs. (H) Internal CD14 MFI for both BMDMs (black) and AMs (pink). (I) Frequency of Q3^+^ (internal CD14^+^) BMDMs and AMs. Data representative of 3 independent experiments (D, G) or compiled from 3 independent experiments (B, C, E, F, H, I). (A - I) *P<0.05, ****P < 0.001, ns, not significant. Two-way ANOVA with Sidak’s multiple comparison test.

To distinguish between differences in total TLR4 protein expression versus localization, we performed intracellular staining for TLR4 in untreated and LPS stimulated conditions.

Despite minimal surface expression, AMs expressed relatively high levels of internal TLR4 protein (**Fig 3D-E**). However, internal TLR4 protein levels were still significantly lower in AMs than BMDMs (**Fig. 3E**). A higher percentage of AMs than BMDMs expressed internal, but not surface TLR4, yet at 4 hours post-LPS stimulation BMDMs had an increase in the percentage of cells expressing only internal TLR4 over untreated (**Fig. 3F**), correlating to the internalization seen in **Fig. 3B**. We also measured CD14 surface and internal expression in AMs and BMDMs. Overall, AMs expressed lower levels of internal CD14 than BMDMs (**Fig 3G-H).** Similar to the trends observed for TLR4 expression, a higher percentage of AMs, compared to BMDMs, expressed internal but not surface CD14 (**Fig. 3I**). Overall, AMs have significantly lower surface expression of TLR4 and CD14 than BMDMs, yet do express some TLR4 and CD14 protein in intracellular stores. This difference in receptor/co-receptor expression and localization likely affects the ability of AMs to sense and respond to low-dose LPS. Our data suggest that even though there is an increase in CD14 surface expression after 4 hours of LPS stimulation in AM (**Fig 3C**), this change is not enough to facilitate LPS sensing at low doses.

### Alveolar macrophages do not produce IL-10 due to low expression of c-Maf

When evaluating AM responses to LPS stimulation, we initially hypothesized that excessive IL-10 production could be an explanation for the AM hyporesponsive state. IL-10 is an immunosuppressive cytokine induced downstream of both TLR and Type I IFN signaling^26,27^. In BMDMs, LPS-induced IL-10 production is partially dependent on the activation of both the TRIF and MYD88 pathways^28^, triggered by endosomal and surface-located TLR4 respectively. IL-10 is also induced through autocrine signaling of Type I IFNs^29^. When we tested AM IL-10 production, we instead observed that AMs produced no detectable IL-10 across all LPS doses tested (0.1 - 10 ng/mL) (**Fig. 4A**). Additionally, we detected little to no IL-10 produced by AMs in response to Pam3Cys, R848, and CpG, PAMPs for which AMs mount robust TNF and IL-6 responses (**Fig. S4A, B, C**). In contrast, BMDMs produced IL-10 in response to LPS, Pam3Cys, R848, and CpG, as expected (**Fig. 4A, S4A, B**).

**Figure 4:**
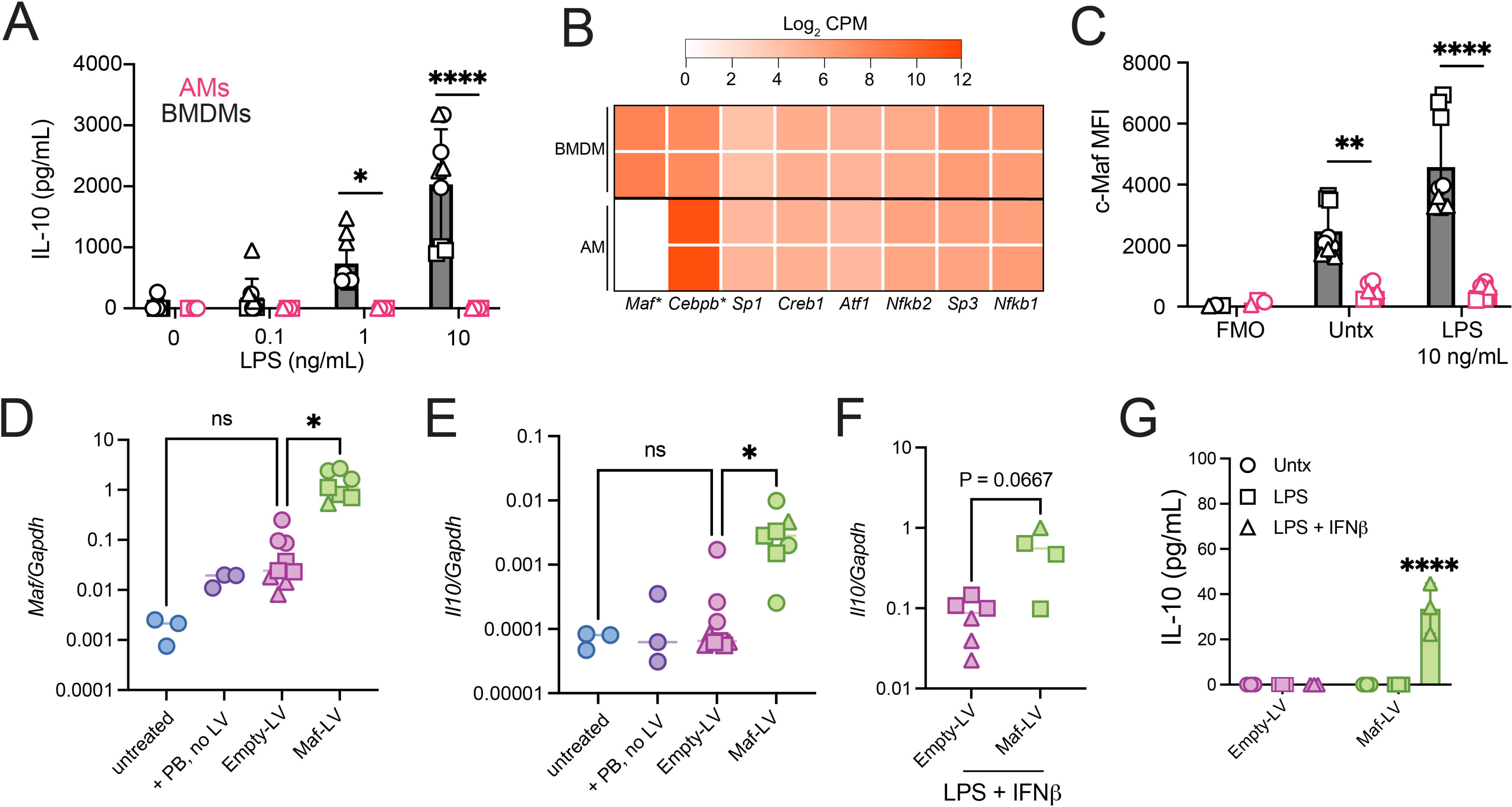
Alveolar macrophages do not produce IL-10 in response to multiple stimuli due to low c-Maf expression. (A) IL-10 for AMs (pink) and BMDMs (black) untreated or stimulated with 0.1, 1, or 10 ng/mL of LPS for 20 hours. (B) Expression of IL-10 promoter-associated genes (log_2_ CPM) from bulk RNA-sequencing of untreated AMs (left) and BMDMs (right) after 4 hours using data from Fig. 1A. *adj p-value < 0.01. (C) c-Maf MFI for BMDMs (black) or AMs (pink) after LPS (10 ng/mL) stimulation or untreated. (D) *Maf* gene expression relative to *Gapdh* for mexAMs treated with media (no polybrene (PB) no virus), PB only, Empty-lentivirus (LV), or Maf-LV. (E) *Il10* gene expression relative to *Gapdh* for mexAMs under the same conditions for (D). (F) *Il10* gene expression relative to *Gapdh* for mexAMs transduced with Empty or Maf LV then stimulated with LPS (10 ng/mL) and rIFNβ (100 ng/mL) for 4 hours. (G) IL-10 for mexAMs transduced with Empty or Maf-LV then stimulated with LPS (10 ng/mL) and IFNβ (100 ng/mL) for 24 hours. Data compiled from 3 independent experiments (A, C-E) or 2 independent experiments (F) or representative of 2 independent experiments (G). (A, C, I) *P<0.05, **P<0.01, ****P<0.0001 by two-way ANOVA with Sidak’s multiple comparison test. (F, G) *P<0.05 by one-way ANOVA with Tukey’s multiple comparisons test. (H) P-value reported by Mann-Whitney test.

Because no IL-10 was detected for by multiple PAMPs, including ones that induced robust TNF and IL-6 responses in AMs (**Fig. S2A,B**), we reasoned that the mechanism must be separate from TLR signaling and proximal to IL-10 expression itself. We examined AM gene expression of transcription factors known to be associated with the IL-10 promoter: *Maf, Atf1, Nfkb1, Nfkb2, Creb1, Cebpb, Sp1,* and *Sp3*^27^. Out of all of the transcription factors, only *Maf* expression, the gene encoding for c-Maf protein, was significantly lower in expression in AMs compared to BMDMs (adj p-value < 0.0001, One-way ANOVA) (**Fig 4B**). Measuring c-Maf protein by flow cytometry, we found that AMs have significantly lower expression of c-Maf compared to BMDMs (**Fig 4C**). This suggested that a lack of c-Maf expression could be preventing AMs from making IL-10.

To test if c-Maf was sufficient to induce IL-10 expression in AMs, we aimed to express c-Maf in AMs exogenously. Manipulation of genes in primary murine AMs is difficult due to their limited growth and lifespan in culture and the initial low yield per animal that limits further alterations. To address the issues of limited lifespan and low yield, we expanded primary AMs isolated from WT mice by culturing them in media containing GM-CSF, TGF-β and PPARL agonist Rosiglitazone, following a published protocol^30^. Murine *ex vivo* AMs, termed “mexAMs”, were shown by Gorki et al. to maintain classic AM surface marker expression through multiple passages^30^. Other studies have shown that primary cultured AMs can maintain AM transcriptional and epigenetic identity when reintroduced to the lung *in vivo*^31^. We first verified that mexAMs do not make c-Maf or IL-10 in response to LPS (**Fig. S4 D,E**). We transduced mexAMs with lentivirus containing either a plasmid with a Maf gene cassette (Maf-LV) or a control plasmid with an empty gene cassette (Empty-LV). After puromycin selection, we detected a significant increase in *Maf* expression in Maf-LV-transduced mexAMs compared to Empty-LV-transduced cells, mexAMs treated with polybrene alone, or untreated mexAMs (**Fig. 4D**). Additionally, baseline *Il10* expression was higher after Maf-LV transduction compared to all other conditions (**Fig. 4E**). After stimulation with LPS and IFNβ, there was a trend towards an increase in *Il10* expression and detection of IL-10 protein in the Maf-LV compared to Empty-LV condition. (**Fig. 4F, G**).

To confirm if *Il10* and *Maf* were also differentially expressed in human AMs, we interrogated single-cell RNA data from the Integrated Cell Atlas of the Lung through the Census database of CZ CELLxGENE Discover^32^, filtering to include AMs, classical monocytes, and lung macrophages within healthy lung tissue. While both classical monocytes and lung macrophages had high expression of *CD14*, human AMs had very low expression (**Fig. S4F**). Similarly, human lung macrophages showed high expression of *MAF*, while both AMs and monocytes had overall low expression (**Fig. S4F**). None of the cell types from the healthy lung showed high expression for *IL10*. *MARCO*, a gene known to have high expression in AM populations, was used as a control. These data suggest that the that AM gene expression levels we have observed in mice for *Il10*, *Maf*, and *Cd14* are reflective of gene expression of AM from healthy human lungs.

### IFN**β** enhances alveolar macrophage TNF and IL-6 response to low-dose LPS in the absence of IL-10

One implication for an absence of IL-10 production by AMs is that conditions that would normally stimulate robust IL-10 production might have alternative impacts. We hypothesized that one of those conditions could be Type I IFN, which has been shown to induce IL-10 in myeloid cells^27^. To test a role for Type I IFN, we stimulated AMs and BMDMs *in vitro* with low-dose LPS (1 ng/mL) and IFNβ (1-100 ng/mL) for 20 hours. BMDMs produced a dose-dependent increase in IL-10 and a significant decrease in TNF and IL-6 with increasing amounts of rIFNβ (**Fig. 5A**). This response recapitulated previously described effects of Type I IFN and IL-10 in macrophages^29^. In contrast, we observed an increase in AM TNF production, peaking at 10 ng/ml IFNβ, and as well as a trending increase in IL-6, with no production of IL-10 (**Fig. 5B**). We also tested AM and BMDM responses to IFNγ and LPS, as IFNγ has been shown to enhance macrophage responses under some conditions^33–35^. In contrast to IFNβ, IFNγ led to an increase in both TNF and IL-6 for both BMDMs and AMs and a significant decrease in IL-10 for BMDMs (**Fig. S5A,B**). To test whether IFNβ altered surface expression of TLR4, we measured TLR4 by flow cytometry after stimulation with rIFNβ and LPS in AM and BMDM. We observed no difference in TLR4 surface in AMs after LPS and IFNβ co-stimulation (**Fig. S5C).** There was also no change in AM c-Maf expression in AMs after 20 hours of incubation with either rIFNβ alone or rIFNβ with LPS (**Fig. S5D**).

**Figure 5:**
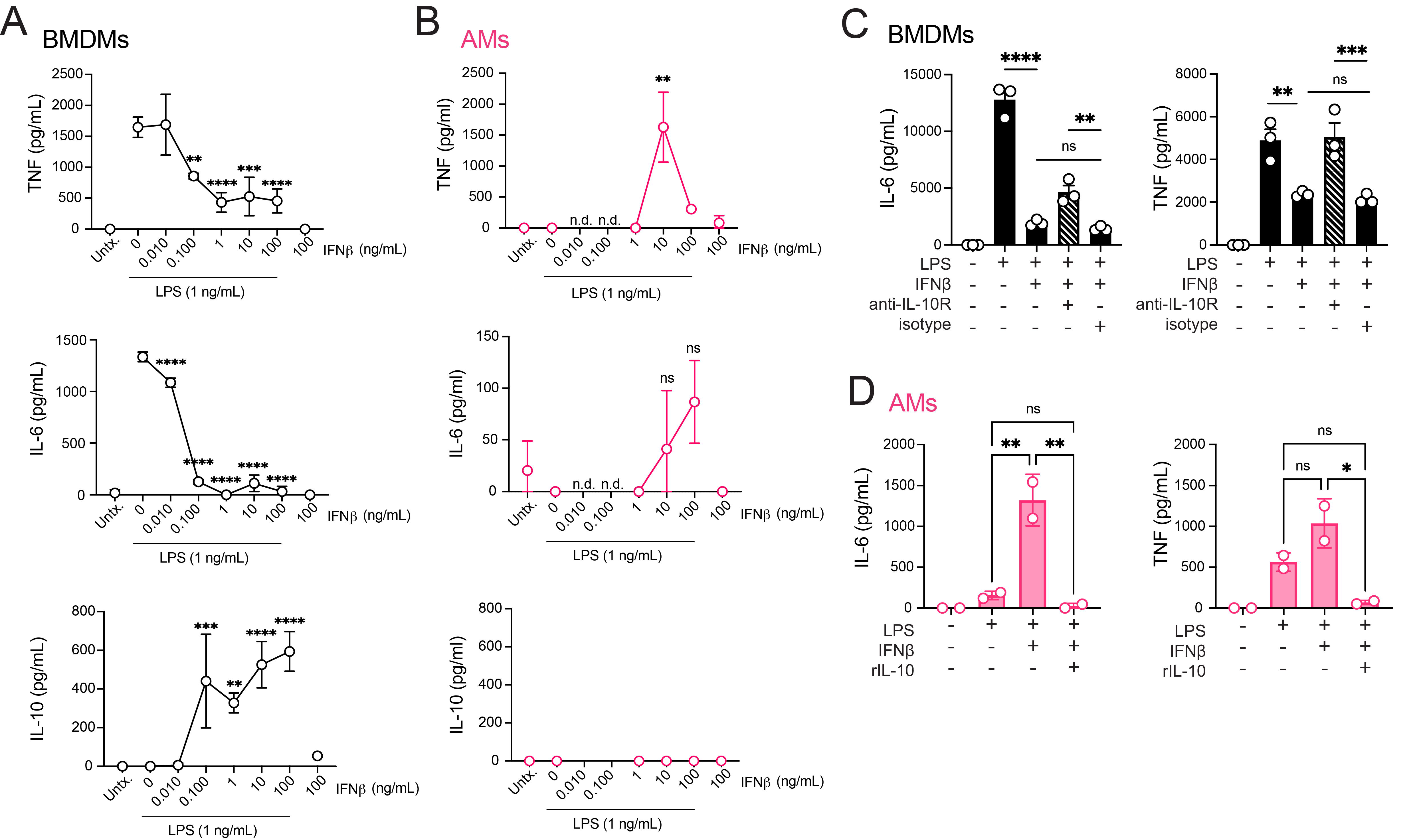
**IFN**β **enhances alveolar macrophage TNF and IL-6 response to low-dose LPS in the absence of IL-10.** (A) TNF, IL-6, and IL-10 for BMDMs stimulated with LPS (1 ng/mL) with 0.01 – 100 ng/mL of rIFNβ. Data shown are mean with SD. with values (B) TNF, IL-6, and IL-10 for AMs stimulated with LPS (1 ng/mL) with 1 - 100 ng/mL of rIFNβ. (C) BMDMs stimulated with LPS (1 ng/mL), rIFNβ (10 ng/mL), anti-IL-10R (10 mg/mL), and/or isotype control (Rat IgG1k, 10 μg/mL). (D) AMs stimulated with LPS (1 ng/mL), rIFNβ (10 ng/mL), and/or rIL-10 (50 mg/mL) for 20 hours followed by ELISA for IL-6 (left) and TNF (right). Data are representative of 3 independent experiments (A-D). **P<0.01, ***P<0.001, ****P<0.0001, n.d. no data, ns not significant. (A, B) One-way ANOVA with Sidak’s multiple comparison test, compared to LPS only condition. (C, D) One-way ANOVA with Tukey’s multiple comparison test.

Next, we aimed to determine the role for IL-10 in the distinct responses to IFNβ between AMs and BMDMs. First, we blocked IL-10 signaling in BMDMs using an anti-IL-10R antibody 24 hours prior to LPS and IFNβ co-stimulation. We observed complete rescue of TNF and partial rescue of IL-6 production in the presence of anti-IL-10R blocking antibody (**Fig. 5C**). Second, we added rIL-10 to AMs stimulated with LPS and IFNβ and found that exogenous IL-10 significantly decreases AM responses (**Fig. 5D**). Overall, these results indicate that an absence of IL-10 production by AMs results in their unique sensitivity to IFNβ and that the enhancement of AM TNF and IL-6 innate responses to LPS by IFNβ is independent of changes in TLR4 surface expression.

## DISCUSSION

AMs serve a critical role as airway sentinels, yet our understanding of AM innate sensing is relatively limited compared to other macrophage subsets, especially in evaluation of direct sensing. In the context of acute lung injury models^35–38^, murine and human AMs produce pro-inflammatory cytokine and chemokines *in vivo* in response to LPS. However, those studies do not assess direct AM cell-intrinsic sensing, as LPS is delivered via the intranasal route to multiple cell types in the airway all at once. This leads to a synchronized innate response across many responding cell types that affect the AM response and is likely very different from the early stages of respiratory infections where there are limited numbers of bacteria. This is the case for an infection such as Mtb. At the early stages of Mtb infection, AMs are the first cells in the lung to become infected, respond in an isolated manner, and fail to generate an inflammatory response within the first days of infection^17,18^. For any lower-respiratory tract infection with a limited initial dose, we predict that AM sensing would have an outsized role on the earliest innate response.

Here, we evaluate AM LPS-specific innate sensing both *in vivo* and *ex vivo* using bead-conjugated and low-dose approaches that enable interrogation of AM direct sensing. We show that AMs produce similar amounts of TNF and IL-6 as BMDMs in response to high concentrations of soluble LPS. However, at lower LPS concentrations or when delivered on coated beads, AMs generate significantly less TNF, IL-6, and Type I IFN signaling than BMDMs. AMs express low levels of TLR4 on the surface yet contain substantial internal pools of TLR4.

Low surface TLR4 in AMs is associated with very low CD14 expression, a co-receptor known to be required for efficient TLR4 trafficking^19^. Based on known roles of CD14 in other sensing pathways, we predict that low CD14 expression may also impact AM’s ability to sense other PAMPs. For example, CD14 has been shown to be required for sensing of Mtb component trehalose 6,6’-dimycolate (TDM) along with TLR2 and MARCO^40^, and other TLR2 associated mycobacterial lipoproteins^41^. CD14 is also known to contribute to the recognition of necrotic cells^42^, several different types of LPS from gram-negative bacteria ^43^, flagellin^44^, and *Legionella pneumophila*^45^. Further investigation into how CD14 expression levels impact AM-specific pathogen sensing will shed light on whether CD14 expression is significantly impacting the early pulmonary response to other respiratory pathogens.

In addition to a role for CD14, it is possible there are other mechanisms that also limit TLR4 trafficking in AMs. One possibility is that AMs have a cell-intrinsic deficiency in one or several components required for endosomal trafficking of TLR4; there are several potential candidates based on previous literature. The GTPase Rab11a is required for recycling of TLR4; minimal Rab11a expression leads to diminished surface expression of TLR4^46,47^. GTPase Arf6 is also required for LPS internalization and trafficking of the adaptor TRAM to the endosome^48^. Additionally, TLR4 internalization requires dynamin^49^. To determine the potential role for these different proteins in regulating TLR4 surface expression, a more extensive screening of GTPases and trafficking regulators is needed for AMs.

While interrogating AM responses to PAMPs, we observed that AMs do not produce IL- 10 in response to any of the stimuli tested. Our findings are supported by a previous study which found that AMs did not produce IL-10 following LPS stimulation and it was independent of calcium entry or intracellular cAMP levels^50^. Additionally, another study found AMs produced no IL-10 *in vivo* after i.p. LPS administration, and have minimal Maf expression by RNA-seq under basal conditions^36^. Here, we show that the absence of IL-10 production in AMs is due to reduced expression of the transcription factor c-Maf. We demonstrate that exogenous expression of c-Maf through lentiviral transduction leads to an increase in IL-10 production by AMs. Interestingly, low expression of Maf-B and c-Maf are associated with self-renewal of macrophages^51,52^, and so we hypothesize that the lack of c-Maf expression we observe in AMs likely enables enhanced proliferative capacity and renewal *in vivo*. A prior study found that Maf expression is increased in AMs during aging and this correlates with a decrease in cell cycling or proliferation^53^. We predict that an increase in Maf expression might also explain why AMs appear to acquire the ability to produce IL-10 during aging^54^. In this way, c-Maf mediates a trade-off in AMs between proliferative capacity and production of IL-10. While production of IL- 10 early during an infection response might be detrimental to mounting effective host immune responses^55,56^, IL-10 is critical during chronic disease and inflammation to prevent immunopathology, including under conditions such as Acute Lung Injury (ALI)^57^. Further investigation is needed into the potential *in vivo* and context-dependent impacts of AM IL-10 production.

We demonstrate that one corollary of the absence of IL-10 production by AMs is their unique response to Type I IFNs. Type I IFNs are critical for anti-viral immunity, but can be detrimental for control of bacterial infections, including enhancing cell death and driving immunopathology^58^. We found that exogenous IFNβ enhanced AM production of TNF and IL-6 in response to low-dose LPS. This is supported by a recent study demonstrating that adenoviral-induced IFNβ enhances AM and AM-like responses to LPS^59^. IFNβ did not appear to directly enhance PRR or adaptor expression and it is still unknown what mediates this enhanced response. It is also unknown how long this IFN-mediated “boost” lasts in AMs and whether IFNβ may generate long-term transcriptional and/or epigenetic remodeling in AMs.

We found that IFNγ also enhances AM IL-6 and TNF responses to LPS in both AMs and BMDMs. IFNγ is commonly used to polarize macrophages to an “M1” inflammatory phenotype^60^ and is associated with host control in both bacterial and viral lung infections. Recent studies have shown that T cell-derived IFNγ is important for induction of MHC II expression in AMs, remodeling of AM responses after viral infection^61,62^ and for enhancing myeloid progenitors from the bone marrow after BCG vaccination^63^. Our data shows that IFNγ enhances an inflammatory response in AMs when coupled with LPS stimulation, but further investigation is warranted to assess the mechanism and the durability of its effects. Future studies to examine the direct effects of Type I and Type II IFNs in AMs are ongoing. Taken together, our results show that AMs have a reduced cell-intrinsic sensing capacity for LPS but are highly sensitive to external IFN signals that allow AMs to mount a more pro-inflammatory response.

## Limitations of the study

By evaluating direct innate sensing, our study uncovers unique cellular mechanisms of AMs that make their innate responses distinct from that of other macrophages. Differences in CD14 expression, TLR4 trafficking, IL-10 production, and sensitivity to Type I IFNs have implications for direct LPS sensing by AMs as well as other innate pathways. However, there are a few limitations of this work that require further investigation. First, we chose to focus on responses to a small selection of PAMPs and on TNF, IL-6, and Type I IFN read-outs of innate signaling, because they represent downstream events for some of the most dominant intracellular pathways. There are many other disease-relevant PAMPs and PRR pathways worth investigating. Second, due to the difficulty in measuring IFNβ protein levels, we were only able to assess differences in AM versus BMDM gene expression for *Ifnb1* and *Ifit3* (IFNβ protein detection using anti-IFN-β ELISA reagents and a reporter cell line were unsuccessful). Third, due to our desire to examine direct sensing by AMs and cell-intrinsic regulation of these pathways, we opted to perform the majority of our studies *in vitro,* rather than *in vivo,* acknowledging the known changes that occur in macrophages following their removal from their microenvironment^31,64–66^. However, we are confident in our *in vitro* approaches to interrogate these pathways, given that the AM responses to LPS sensing, c-Maf expression, and IL-10 production were detected both *in vivo* and *in vitro*. In the future, we plan to follow up on many of these pathways including CD14 expression, TLR4 trafficking, and IL-10 production to assess their *in vivo* relevance during disease. Fourth, as mentioned above, we predict that both Type I and Type II IFN exposures regulate macrophage responses beyond simply IL-10 production and enhancement of IL-6 and TNF production. There are many additional mechanistic studies to perform to fully understand their effects during health and disease.

## STAR METHODS

### EXPERIMENTAL MODEL AND SUBJECT DETAILS

#### Mice

C57BL/6J mice (000664) were purchased from the Jackson Laboratory (Bar Harbor, ME) and bred and maintained in specific pathogen-free conditions under a controlled day-night cycle and given food and water *ad libitum*. 6- to 12-week old male and female mice were used for all experiments except for RNA-Seq which used only female mice. Animal studies for transcriptional analysis of AM and BMDM responses to LPS-coated bead were performed at Seattle Children’s Research Institute in compliance with and approval by the Seattle Children’s Research Institute’s Institutional Animal Care and Use Committee. All other animal studies were performed at University of Massachusetts Amherst in compliance with and approval by the University of Massachusetts Amherst’s Institutional Animal Care and Use Committee.

#### Primary cells

For mouse alveolar macrophage (AM) preparation, bronchoalveolar lavage was performed by first exposing and puncturing the trachea of euthanized mice with Vannas Micro Scissors (VWR, 76457-352). 1 mL of cold PBS (gibco, 10010-049) was injected into the lungs with a 20-gauge IV catheter (Braun, 4252543-02) with a 1 mL syringe (McKesson, 16-PS1C) and the lungs were flushed a total of 4 times. For each wash, the PBS was collected into a conical tube over ice, filtered (70 μm, Falcon, 352350) then spun down and counted. Cells were plated at a density of 0.5 – 1 * 10^5 cells per well in a 96-well plate, in RPMI (Millipore Sigma, 11875-119) with 10% FBS (Biowest, S1620) with 1% L-glutamine (2 μM, gibco, 25030-081) and 1% penicillin-streptomycin (100 U/mL, gibco, 15140-122). AMs adhered overnight at 37°C with 5% CO_2_ before experiments were conducted. Each experiment replicate included BAL pooled from 5-10 mice (∼1.5 mice per well).

For bone-marrow derived macrophages (BMDM), bone marrow was isolated from murine femurs by flushing the bone with RPMI. Cells were filtered, spun down, and plated on 15 cm non-TC treated plates (VWR, 25384-326) and cultured for 6 days in RPMI with 10% FBS, 1% L-glutamine, 1% penicillin-streptomycin, and 0.01% recombinant human M-CSF (0.05 μg/mL, Peprotech, 300-25). For experiments using ELISA and RT-qPCR, cells were replated at the same confluency as collected AMs in a 96-well plate. For other experiments including flow cytometry or ELISA independent of AMs, cells were plated at 0.25 – 1 * 10^6 cells per well in a 24-well or 12-well plate. After replating, BMDMs adhered overnight at 37°C, 5% CO_2_ before stimulating conditions were added.

#### Cell culture

293T cells (ATCC, CRL-3216) were maintained in Dulbecco modified essential medium (DMEM) (gibco, 11965-092) containing 10% FBS, 1% penicillin/streptomycin, 1% L-glutamine, 1% HEPES (gibco, 15630-080), 1% sodium pyruvate (gibco, 11360-070), and 1% MEM amino acid solution (gibco, 11130-051) at 37°C, 5% CO_2_ in a humidified incubator.

### METHOD DETAILS

#### Generation of LPS Beads

For RNA-sequencing experiments, Polysciences Fluoresbrite Carboxylate Microspheres 1.00 um (YG) were incubated overnight with 100 µg/mL LPS (R595 S. Minnesota) in PBS or PBS alone at 4 degrees. The next day, the beads were washed 10X in PBS. Each wash was followed by centrifugation at 10,000 x g for 5 minutes. Beads were then resuspended in PBS. For flow cytometry experiments, stock solution of Streptavidin Fluoresbrite® YG Microspheres (1.0 μm, yellow-green fluorescent, Polysciences, 24161-1) were vortexed and 3 μL were mixed with 1.5 μg of LPS-EB Biotin (InvivoGen, tlrl-lpsbiot) and incubated overnight at 4°C, covered. The next day, 100 μL of sterile 1% BSA in PBS was added to the bead mixture, mixed, then centrifuged at 10,000 x *g* for 5 min. The BSA PBS was carefully aspirated with a pipette and the wash was repeated a total of three times. After the final wash, the beads were resuspended in 1 mL of 1% BSA in PBS.

#### Bead aerosolization

Beads generated were aerosolized using a LC Sprint^®^ Reusable Nebulizer (PARI, 023F35) attached to a mouse cage with a vacuum pump and air flow regulator. Beads were resuspended in 4 mL of ddH2O and delivered at 3 liters/minute for 20 minutes. Treated mice were rested for 4 hours (for RNA-sequencing) or 30 minutes (for 20 hour Brefeldin A incubation and flow cytometry) prior to euthanasia.

#### RNA-Seq and Analysis

RNA isolation was performed using TRIzol (Invitrogen, 15596018), two sequential chloroform extractions, Glycoblue carrier (Invitrogen, AM9515), isopropanol precipitation, and washes with 75% ethanol. RNA was quantified with the Bioanalyzer RNA 6000 Pico Kit (Agilent, 5067-1513). cDNA libraries were constructed using the SMART-Seq v4 Ultra Low Input RNA Kit (TaKaRa, 634889) following the manufacturer’s instructions. Libraries were amplified and then sequenced on an Illumina NovaSeq 6000 (150bp paired-end). The read pairs were aligned to the mouse genome (mm10) using the gsnap aligner^67^. Concordantly mapping read pairs (∼20 million / sample) that aligned uniquely were assigned to exons using the 25iocond program and gene definitions from Ensembl Mus_Musculus GRCm38.78 coding and non-coding genes. Genes with low expression were filtered using the “filterByExpr” function in the edgeR package^68^.

Differential expression was calculated using the “edgeR” package. Heat map visualizations were generated in R using the ‘heatmap.2’ library.

#### Gene Set Enrichment Analysis (GSEA)

Input data for GSEA consisted of lists, ranked by -log(p-value), comparing RNAseq expression measures of target samples and controls including directionality of fold-change. Mouse orthologs of human Hallmark genes were defined using a list provided by Molecular Signatures Database (MsigDB)^69^. GSEA software was used to calculate enrichment of ranked lists in each of the respective hallmark gene lists, as described previously^70^. A nominal p-value for each ES is calculated based on the null distribution of 1,000 random permutations. To correct for multiple hypothesis testing, a normalized enrichment score (NES) is calculated that corrects the ES based on the null distribution. A false-discovery rate (FDR) is calculated for each NES.

#### *Ex vivo* bead addition with Brefeldin A

After cells were isolated and let adhere overnight, beads were added at an optimized concentration where 5-10% of cells were “Bead+” as identified by flow cytometry. For LPS Beads, 2.5 μL of bead suspension was resuspended in 1 mL of appropriate cell culture media, and 100 μL was deposited onto plated cells in a 96-well plate. Cells were concurrently given 100 μL of Brefeldin A (1000X solution diluted to 2X, BioLegend, 420601) to arrest protein secretion to measure cytokines via intracellular staining. Cells were stimulated with beads in the presence of Brefeldin A for 20 hours prior to analysis.

#### Quantitative reverse transcription PCR (RT-qPCR)

Cells were plated at a density of 0.5 – 1 * 10^5 cells per well in a 96-well plate, followed by stimulation. RNA isolation was performed using TRIzol (Invitrogen, 15596018), two sequential chloroform extractions, Glycoblue carrier (Invitrogen, AM9515), isopropanol precipitation, and washes with 75% ethanol. RNA was quantified with the Biodrop Duo (Biochrom). Equivalent amounts of RNA (1 μg per sample) were converted to complementary DNA (cDNA) and amplified using RNA to cDNA EcoDry Premix (TaKaRa, 639543) per the manufacturer’s instructions. RT-qPCR was performed using TaqMan primer probes (IDT) with TaqMan Fast Universal PCR Master Mix (Applied Biosystems, 4352046) using a BioRad CFX Opus 96 RT-qPCR detection system. Data was normalized to relative *Gapdh* expression in individual samples.

#### Enzyme-linked immunosorbent assay (ELISA)

Cells were plated either at a density of 0.5 – 1 * 10^5 cells per well in a 96-well plate for experiments involving both AMs and BMDMs, or 0.25 – 1 * 10^6 cells per well in a 24-well or 12- well plate for experiments with BMDMs alone. After 20 hours of stimulation, supernatant was removed from individual wells and stored at -20°C overnight. Murine IL-6 (DY406), TNF (DY410), and IL-10 (DY417) DuoSet enzyme-linked immunosorbent assays (ELISAs) were performed per manufacturer’s instructions (R&D Systems).

#### Flow cytometry intracellular staining

Cells were collected into sterile tubes and subsequently stained with Fc block (Biolegend, 101320), Live/Dead stain (Zombie Violet Fixable Viability Kit, Biolegend) and characteristic cell surface markers (AMs, Siglec-F (anti-mouse CD170, clone S17007L, Biolegend), CD11c (clone N148, Biolegend); BMDMs F4/80 (clone BM8, Biolegend)). Cells were resuspended in 200 μL of Cyto-Fast™ Fix/Perm Buffer (Biolegend, 426803) and incubated for 20 minutes at room temperature and washed with Cyto-Fast™ Perm Wash (Biolegend, 426803). TNF and IL-6 antibodies were diluted 1:100 in Cyto-Fast™ Perm Wash and added to the cells for 20 minutes. Cells were then washed and fixed with 2% PFA (Electron Microscopy Sciences, 15713-S) prior to acquisition. For phospho-STAT1 staining (clone A15158B, Biolegend), cells were fixed in 4% PFA for 15 minutes, washed with FACS buffer (PBS, 1% BSA, 0.01% NaN_3_), then resuspended in 200 μL of True-Phos™ Perm Buffer (Biolegend, 425401). Cells were incubated for 1 hour at - 20°C. Cells were then washed with FACS buffer and stained with phospho-STAT1 antibody for 30 minutes at room temperature. Cells were washed once more and resuspended in 200 μL of FACS buffer prior to acquisition. For c-Maf staining, cells were surface stained and resuspended in Foxp3 Fixation/Permeabilization solution (Invitrogen, 00-5523-00) and incubated at 4°C for 30 minutes. Cells were washed twice in Permeabilization Buffer (Invitrogen, 00-5523-00) by centrifuging samples at 400 x *g* for 5 minutes at room temperature. Cells were resuspended in 100 μL of Permeabilization Buffer and 0.5 μg of c-Maf antibody (clone sym0F1, Invitrogen) was added. After 30 minutes, cells were washed twice and resuspended in FACS buffer for acquisition.

#### Maf Lentivirus Generation

Maf (pLenti-GIII-CMV) and Empty (pLenti-III-Blank) lentiviral vectors (ABM) were transformed into ProClone Competent DH5α cells (ABM, E003) by heat-shock. Cells were recovered for 1 hour in 150 μL of sterile LB broth in an incubated shaker set at 37°C, 240 rpm before spreading the entire volume on LB agar plates containing kanamycin (Sigma-Aldrich, L0543). Single colonies were inoculated in 4 mL of LB broth (Fisher, 244620) + kanamycin (Sigma Aldrich, K0254) and incubated for 16 hours. 1 mL of the resulting culture was then added to 99 mL of LB broth + kanamycin and incubated overnight. The overnight bacterial culture was harvested and DNA was eluted via MAXI-prep (QIAGEN, 12162). 293T cells were plated the day before lentivirus packaging on plates coated with poly-L-lysine (Sigma-Aldrich, A-005-C) with 10 million cells per 15 cm TC-treated plate. Lentivirus was packaged using CMV-VSV-G envelope plasmid (Addgene, 8454) and psPax2 packaging plasmid (Addgene, 12260) alongside either Maf or Empty cassettes with polyethylenimine (Polysciences, 24765-100). Transfected cells were incubated overnight and washed the next day. Two days after the wash, lentivirus was isolated and purified by ultracentrifugation and resuspended in PBS and frozen at -80°C prior to use.

#### Human Lung Data Acquisition and Analysis from CZ CELLxGENE: Discover

Publicly-available single cell RNA data from CZ CELLxGENE Discover was accessed via Python (v3.11.7) using the cellxgene_census module. Census data was filtered to include cell types “alveolar macrophage”, “classical monocyte”, and “lung macrophage” with a disease state of “normal” in lung tissue in Homo sapiens, and the genes *Cd14* (ENSG00000170458), *Il10* (ENSG00000136634), *Maf* (ENSG00000178573), and *Marco* (ENSG00000019169) as a positive control for AMs. The data slice was stored as an AnnData object^71^ and saved as an H5AD file. The data slice included 248,180 AMs, 99,390 classical monocytes, and 1,864 lung macrophages from the Census version 2023-12-15. A stacked violin plot was generated to visualize gene expression across selected cell types. The analysis involved normalization to counts per million (CPM) and log transformation of the data, the violin plot was created using SCANPY stacked_violin function^72^.

#### Statistical analyses

Data were analyzed for comparison of multiple comparisons between BMDMs and AMs by two-way analysis of variance (ANOVA) (95% confidence interval) with Sidak’s multiple comparisons test. For comparisons within one cell type, data were analyzed by one-way analysis of variance (ANOVA) (95% confidence interval) with Sidaks’s or Tukey’s multiple comparisons test, as reported. Significance is denoted as: \**P* < 0.05, \*\**P* < 0.01, \*\*\**P* < 0.001, \*\*\*\**P* < 0.0001.

Statistical analysis and graphical representation of data were performed using either GraphPad Prism v10.0 software or R.

## Supporting information

Key Resources Table

## Acknowledgments

We thank the Animal Care Staff at University of Massachusetts Amherst and Seattle Children’s Research Institute. We thank Amy Burnside and the Flow Cytometry Core at the University of Massachusetts Amherst. We thank members of the Rothchild and Pobezinsky labs for helpful discussions.

## Author contributions

Conceptualization: PNL, ACR

Methodology: PNL, ACR, LKP, MMC, SD, DM, AD

Investigation: PNL, ACR, LKP, MMC, AT, DD, DM Visualization: PNL, ACR

Data curation: PNL, ACR, AD Formal analysis: PNL, ACR Project administration: ACR Funding acquisition: ACR, AA, AD Supervision: ACR

Writing – original draft: PNL

Writing – review & editing: ACR, MMC, LKP, AD

## Data and materials availability

Raw and processed RNA-sequencing data can be accessed from the National Center for Biotechnology Information (NCBI) Gene Expression Omnibus (GEO) database under accession number GSExxx (*Submission currently private*).

## SUPPLEMENTARY FIGURE LEGENDS

**Figure S1: Gating strategy for *ex vivo* bead-treated cells and peritoneal macrophage bead response.** A-B) Gating strategy for AMs (A) and BMDMs (B) for cell sorting and collection for RNA-sequencing. C-D) Gating scheme for BMDMs (C) and AMs (D) treated *ex vivo* with LPS-coated beads. E) Percent of Bead+ cells for AMs and BMDMs, including both PBS and LPS Bead conditions. F) TNF and IL-6 ICS of peritoneal macrophages treated *ex vivo* with LPS beads for 20 hours. Data is compiled from 3 independent experiments (E) or two independent experiments (F). Technical replicates within each experiment represented by unique shapes. **P < 0.01, ****P < 0.0001. One-way ANOVA with Tukey’s multiple comparisons test.

**Figure S2: Alveolar macrophage pro-inflammatory response is PAMP and cell-specific**. (A) TNF (left) and IL-6 (right) of BMDMs and AMs stimulated with Pam3Cys (0 - 1000 ng/mL) for 20 hours. (B) TNF (left) and IL-6 (right) of BMDMs and AMs stimulated with R848 (0 - 100 mg/mL) for 20 hours. (C) TNF (left) and IL-6 (right) of PMs stimulated with LPS (0 - 10 ng/mL) for 20 hours. (D) TNF (left) and IL-6 (right) of PMs stimulated with Pam3Cys (0 - 1000 ng/mL) for 20 hours. (E) RT-qPCR of *Ifnb* (left) and *Ifit3* (right) expression relative to Gapdh for 2’3’- cGAMP (0-10 mg/mL) after 4 hours. AMs (pink), technical duplicate; BMDMs (black)/PMs (purple), technical triplicate. Data representative of three independent experiments (A) or two independent experiments (B-D). *P < 0.05, **P < 0.01, ***P < 0.001, ****P < 0.0001. Two-way ANOVA with Sidak’s multiple comparison test.

**Figure S3: Peritoneal macrophage expression of TLR4/CD14 and alveolar macrophage and bone marrow derived macrophage expression of TLR2.** A) FMO staining controls for TLR4 internal and surface and B) CD14 internal and surface staining of BMDMs (top) and AMs (bottom) C) Surface TLR4 expression in PMs after 0, 1, or 4 hours of stimulation with LPS (1 ng/mL). D) CD14 MFI of PMs after 0, 1, or 4 hours of stimulation with LPS (1 ng/mL). E) TLR2 MFI of AMs and BMDMs. Data representative of 2 independent experiments (C-E). **P < 0.01, ***P < 0.001. (C, D) One-way ANOVA with Tukey’s multiple comparisons test, compared to untreated. (E) One-way ANOVA with Tukey’s multiple comparisons test.

**Figure S4: Macrophage IL-10 production and gene expression.** (A) IL-10 for AMs and BMDMs after 20 hours of Pam3Cys (0 - 1000 ng/mL). (B) IL-10 for AMs and BMDMs after 20 hours of R848 (0 - 100 mg/mL). (C) IL-10 for AMs and BMDMs after 20 hours of CpG (0 - 10 mM). (D) c-Maf MFI in untreated mexAMs or after 1 ng/mL of LPS for 20 hours. (E) IL-10 measured by ELISA of mexAMs simulated with 0 - 10 ng/mL LPS for 20 hours. (F) Data acquired from CZ CELLxGENE Discover from healthy human lung tissue. Data shown are raw counts normalized to counts per million and log-transformed. (A-C, E) Data is representative of two independent experiments. **P < 0.01, ***P < 0.001, ****P < 0.0001, Two-way ANOVA with Sidak’s multiple comparison test.

**Figure S5: Macrophage response to LPS and IFN**γ **stimulation, and TLR4 and c-Maf MFI after LPS and IFN**β **stimulation.** (A) TNF, IL-6, and IL-10 for AMs stimulated with LPS (1 ng/mL) and/or rIFNγ (2 - 200 ng/mL) for 20 hours. (B) BMDMs under the same conditions and readout as (A). (C) TLR4 MFI of AMs (pink, left) and BMDMs (black, right) untreated or stimulated with LPS (1 ng/mL) and/or rIFNβ (10 ng/mL) for 20 hours. (D) c-Maf MFI of AMs (pink, left) and BMDMs (black, right) untreated or stimulated with LPS (1 ng/mL) and/or rIFNβ (10 ng/mL) for 20 hours. Data representative of two independent experiments *P<0.05, ***P < 0.001, ****P<0.0001, One-way ANOVA with Sidak’s multiple comparison test.

