## Supplemental Figures 1-5 for "Absence of c-Maf and IL-10 enables Type I IFN enhancement of innate responses to low-dose LPS in alveolar macrophages"

**Supplementary Materials**

Fig. S1: Gating strategy for *ex vivo* bead-treated cells and peritoneal macrophage bead response.

Fig. S2: Alveolar macrophage pro-inflammatory response is PAMP and cell-specific.

Fig. S3: Peritoneal macrophage expression of TLR4/CD14 and alveolar macrophage and bone marrow derived macrophage expression of TLR2.

Fig. S4: Macrophage IL-10 production and gene expression.

Fig. S5: Macrophage response to LPS and IFNγ stimulation, and TLR4 and c-Maf MFI after LPS and IFNβ stimulation.

Table S1: RNA-Sequencing data for *ex vivo* stimulated alveolar macrophages and bone marrow-derived macrophages 4 hours following bead exposure

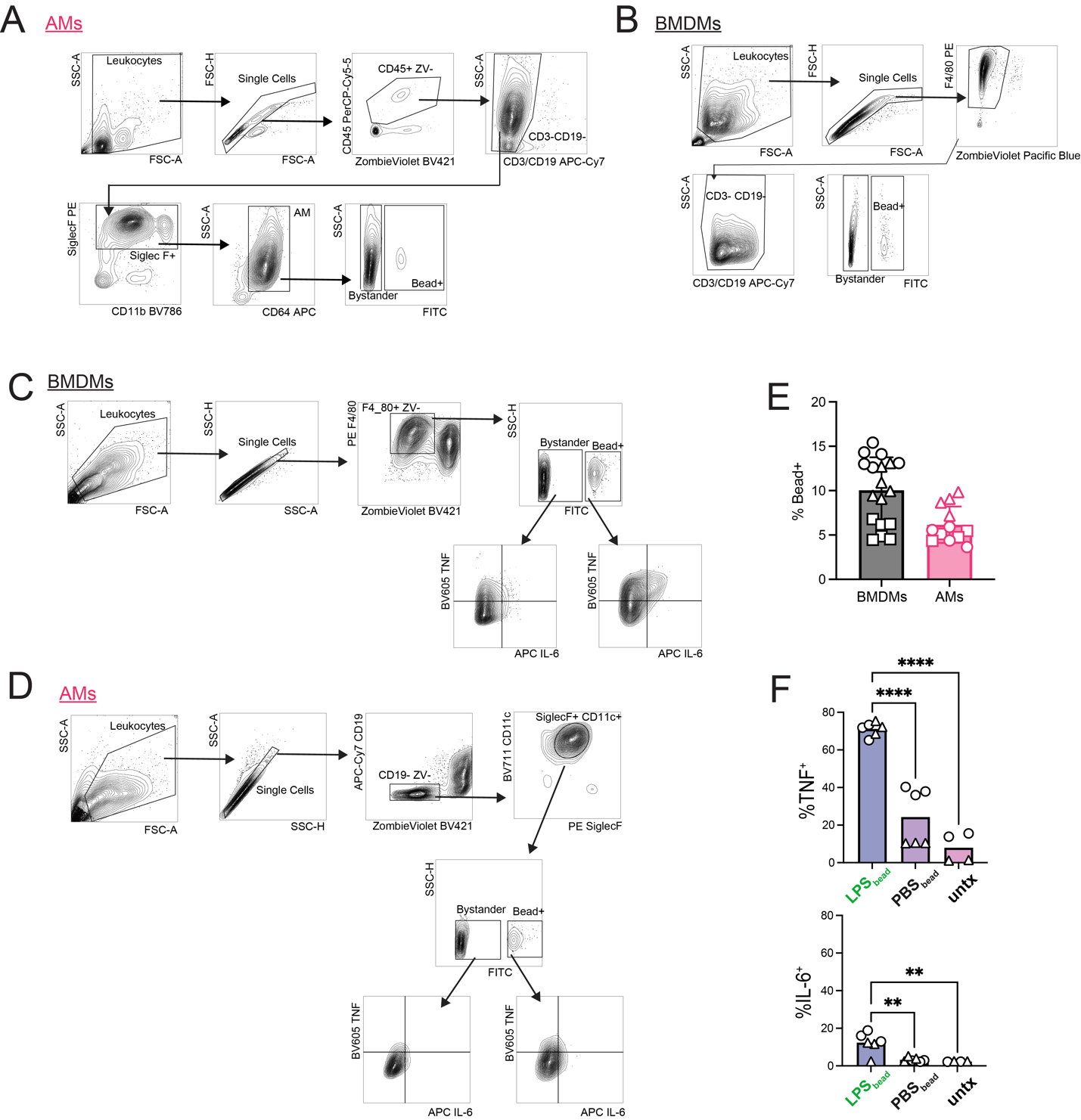

**Figure S1: Gating strategy for *ex vivo* bead-treated cells and peritoneal macrophage bead response.** A-B) Gating strategy for AMs (A) and BMDMs (B) for cell sorting and collection for RNA-sequencing. C-D) Gating scheme for BMDMs (C) and AMs (D) treated *ex vivo* with LPS-coated beads. E) Percent of Bead+ cells for AMs and BMDMs, including both PBS and LPS Bead conditions. F) TNF and IL-6 ICS of peritoneal macrophages treated *ex vivo* with LPS beads for 20 hours. Data is compiled from 3 independent experiments (E) or two independent experiments (F). Technical replicates within each experiment represented by unique shapes. **P < 0.01, ****P < 0.0001. One-way ANOVA with Tukey’s multiple comparisons test.

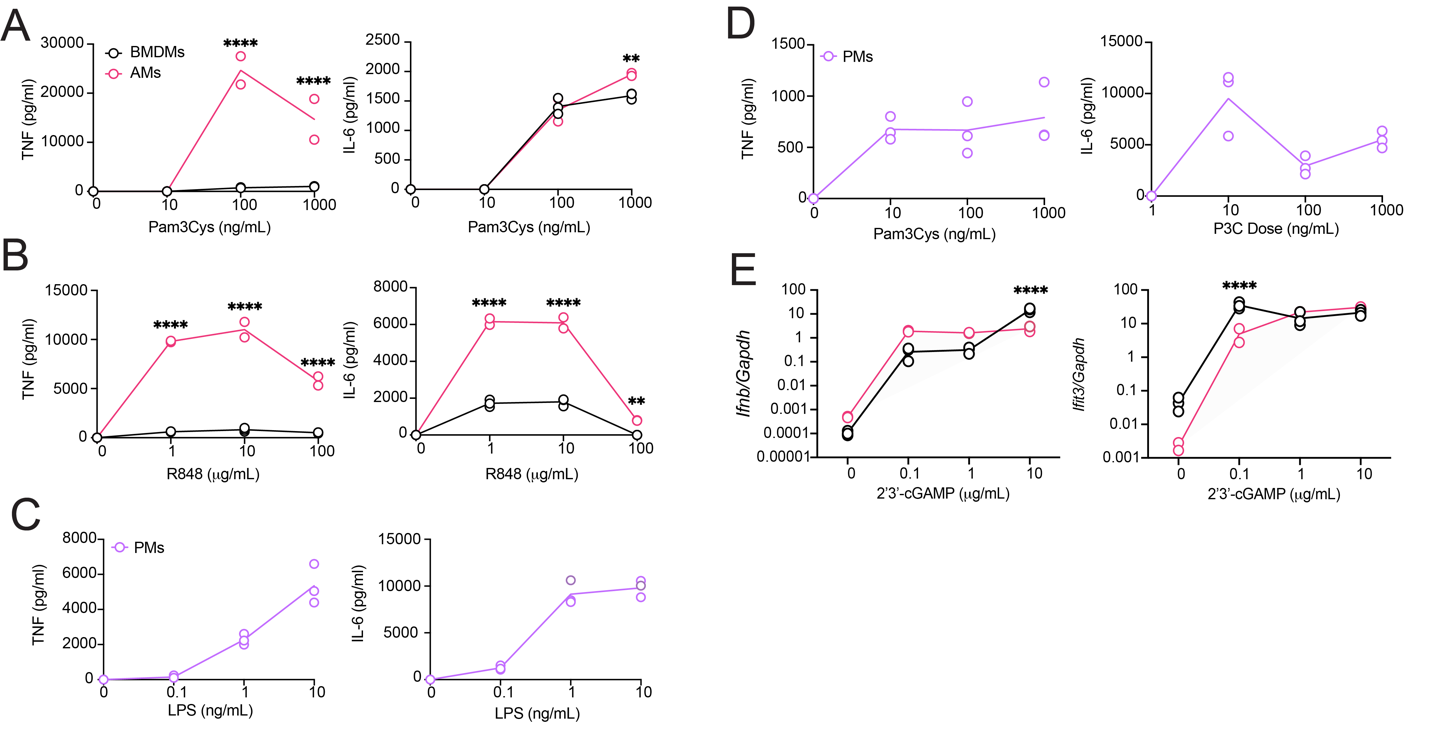

**Figure S2: Alveolar macrophage pro-inflammatory response is PAMP and cell-specific.** (A) TNF (left) and IL-6 (right) of BMDMs and AMs stimulated with Pam3Cys (0 - 1000 ng/mL) for 20 hours. (B) TNF (left) and IL-6 (right) of BMDMs and AMs stimulated with R848 (0 - 100 mg/mL) for 20 hours. (C) TNF (left) and IL-6 (right) of PMs stimulated with LPS (0 - 10 ng/mL) for 20 hours. (D) TNF (left) and IL-6 (right) of PMs stimulated with Pam3Cys (0 - 1000 ng/mL) for 20 hours. (E) RT-qPCR of *Ifnb* (left) and *Ifit3* (right) expression relative to Gapdh for 2’3’-cGAMP (0-10 mg/mL) after 4 hours. AMs (pink), technical duplicate; BMDMs (black)/PMs (purple), technical triplicate. Data representative of three independent experiments (A) or two independent experiments (B-D). *P < 0.05, **P < 0.01, ***P < 0.001, ****P < 0.0001. Two-way ANOVA with Sidak’s multiple comparison test.

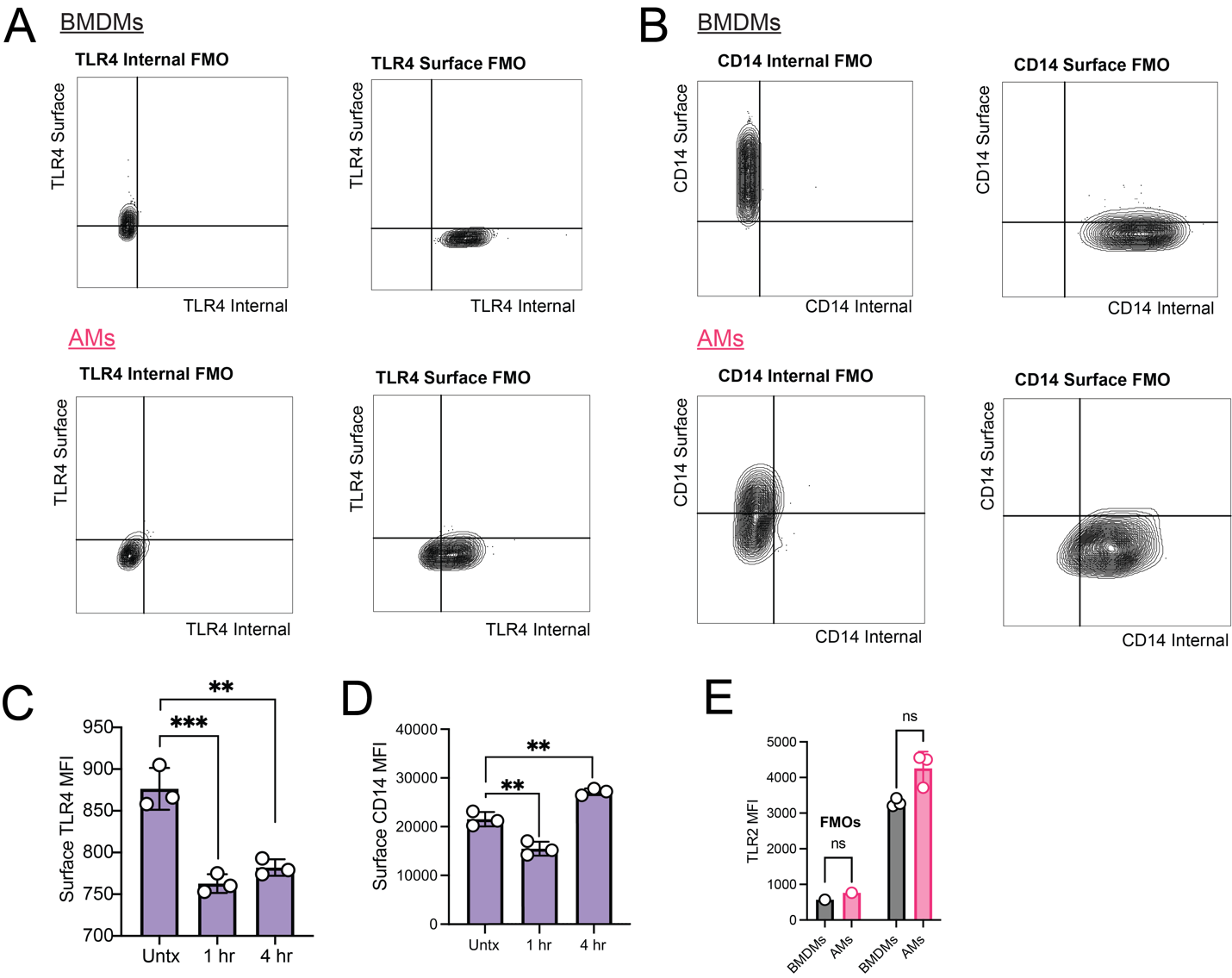

**Figure S3: Peritoneal macrophage expression of TLR4/CD14 and alveolar macrophage and bone marrow derived macrophage expression of TLR2.** A) FMO staining controls for TLR4 internal and surface and B) CD14 internal and surface staining of BMDMs (top) and AMs (bottom) C) Surface TLR4 expression in PMs after 0, 1, or 4 hours of stimulation with LPS (1 ng/mL). D) CD14 MFI of PMs after 0, 1, or 4 hours of stimulation with LPS (1 ng/mL). E) TLR2 MFI of AMs and BMDMs. Data representative of 2 independent experiments (C-E). **P < 0.01, ***P < 0.001. (C, D) One-way ANOVA with Tukey’s multiple comparisons test, compared to untreated. (E) One-way ANOVA with Tukey’s multiple comparisons test.

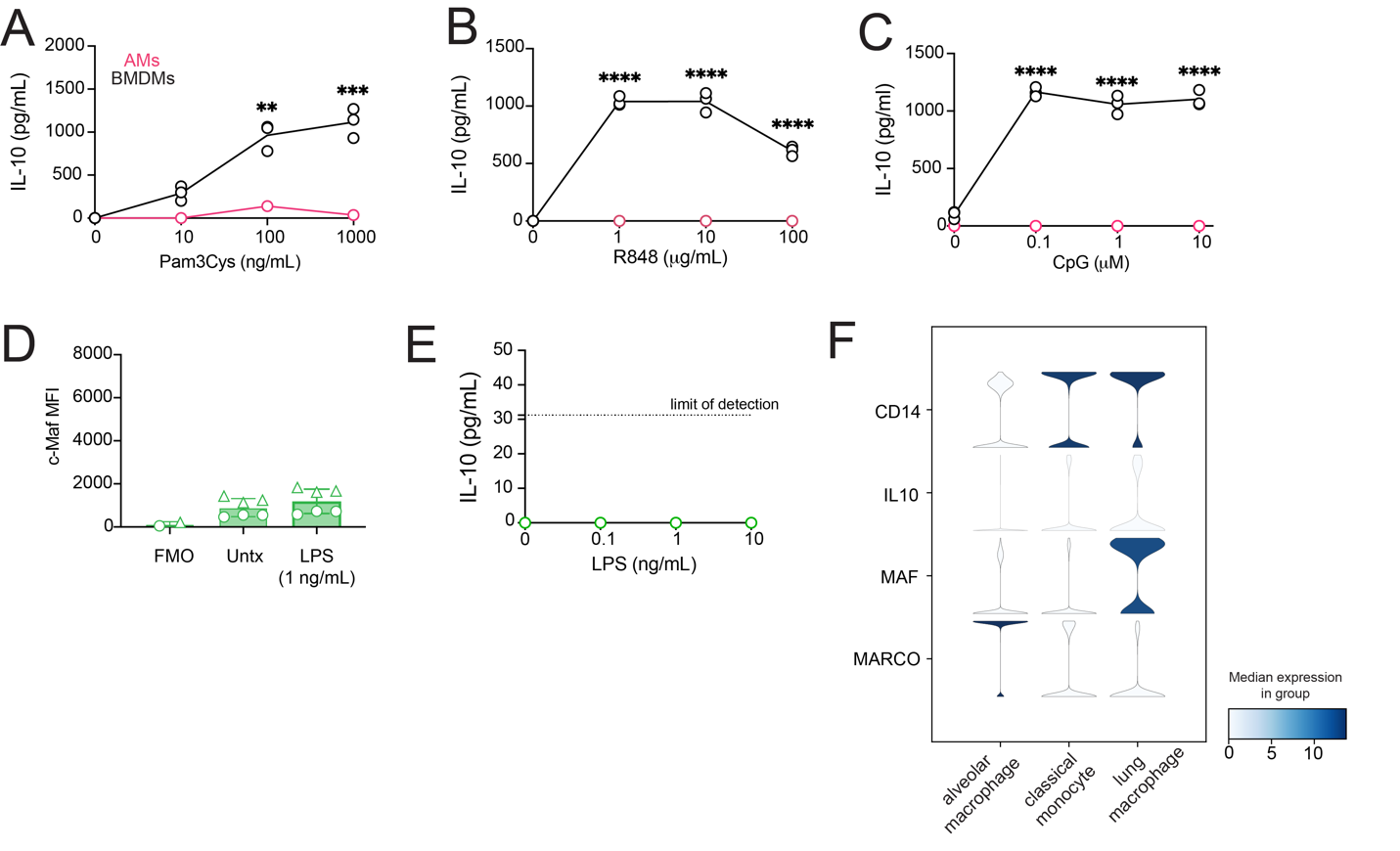

**Figure S4: Macrophage IL-10 production and gene expression.** (A) IL-10 for AMs and BMDMs after 20 hours of Pam3Cys (0 - 1000 ng/mL). (B) IL-10 for AMs and BMDMs after 20 hours of R848 (0 - 100 mg/mL). (C) IL-10 for AMs and BMDMs after 20 hours of CpG (0 - 10 mM). (D) c-Maf MFI in untreated mexAMs or after 1 ng/mL of LPS for 20 hours. (E) IL-10 measured by ELISA of mexAMs simulated with 0 - 10 ng/mL LPS for 20 hours. (F) Data acquired from CZ CELLxGENE Discover from healthy human lung tissue. Data shown are raw counts normalized to counts per million and log-transformed. (A-C, E) Data is representative of two independent experiments. **P < 0.01, ***P < 0.001, ****P < 0.0001, Two-way ANOVA with Sidak’s multiple comparison test.

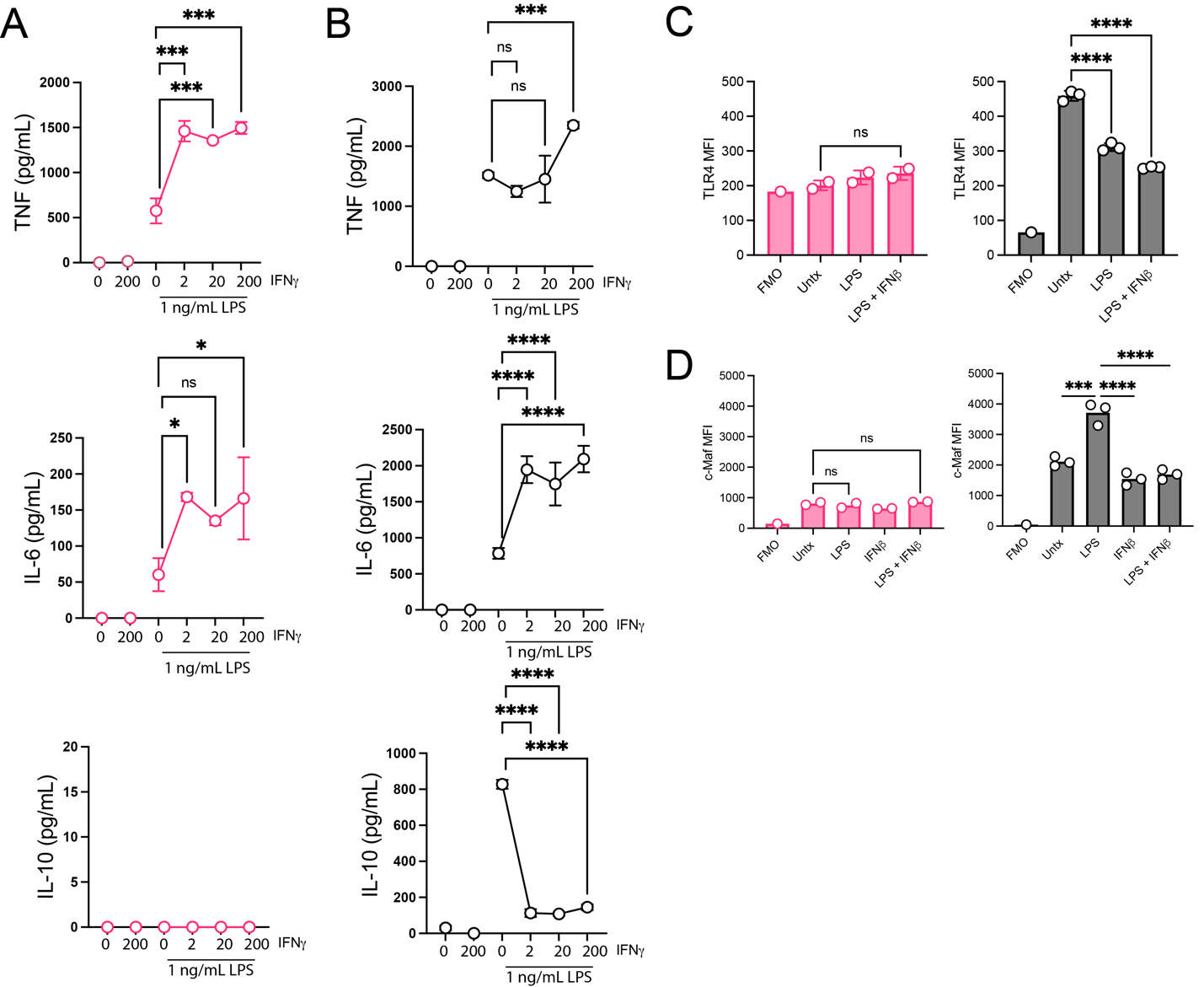

**Figure S5: Macrophage response to LPS and IFNγ stimulation, and TLR4 and c-Maf MFI after LPS and IFNβ stimulation.** (A) TNF, IL-6, and IL-10 for AMs stimulated with LPS (1 ng/mL) and/or rIFN**γ** ( 2 - 200 ng/mL) for 20 hours. (B) BMDMs under the same conditions and readout as (A). (C) TLR4 MFI of AMs (pink, left) and BMDMs (black, right) untreated or stimulated with LPS (1 ng/mL) and/or rIFNβ (10 ng/mL) for 20 hours. (D) c-Maf MFI of AMs (pink, left) and BMDMs (black, right) untreated or stimulated with LPS (1 ng/mL) and/or rIFNβ (10 ng/mL) for 20 hours. Data representative of two independent experiments. *P<0.05, ***P < 0.001, ****P<0.0001, One-way ANOVA with Sidak’s multiple comparison test.
