## Supplementary material for "Absence of c-Maf and IL-10 enables Type I IFN enhancement of innate responses to low-dose LPS in alveolar macrophages": Key Resources Table

| REAGENT or RESOURCE | SOURCE | IDENTIFIER |
| --- | --- | --- |
| Antibodies | | |
| FITC anti-mouse CD45 Antibody (clone 30-F11) | BioLegend | Cat#: 103108 RRID: AB_312973 |
| PE anti-mouse CD45 Antibody (clone 30-F11) | BioLegend | Cat#: 103106 RRID: AB_312971 |
| PerCP/Cyanine5.5 anti-mouse CD45.2 Antibody (clone 104) | BioLegend | Cat#: 109828 RRID: AB_893350 |
| PE/Cyanine7 anti-mouse CD45 Antibody (clone 30-F11) | BioLegend | Cat#: 103114 RRID: AB_312979 |
| APC anti-mouse CD45 Antibody (clone 30-F11) | BioLegend | Cat#: 103112 RRID: AB_312977 |
| APC/Fire™ 750 anti-mouse CD45 Antibody (clone I3/2.3) | BioLegend | Cat#: 147714 RRID: AB_2750441 |
| Brilliant Violet 421™ anti-mouse CD45.2 Antibody (clone 104) | BioLegend | Cat#: 109832 RRID: AB_2565511 |
| Brilliant Violet 510™ anti-mouse CD45 Antibody (clone 30-F11) | BioLegend | Cat#: 103138 RRID: AB_2563061 |
| Brilliant Violet 605™ anti-mouse CD45 Antibody (clone 30-F11) | BioLegend | Cat#: 103139 RRID: AB_2562341 |
| Brilliant Violet 711™ anti-mouse CD45 Antibody (clone 30-F11) | BioLegend | Cat#: 103147 RRID: AB_2564383 |
| Brilliant Violet 785™ anti-mouse CD45 Antibody (clone 30-F11) | BioLegend | Cat#: 103149 RRID: AB_2564590 |
| PE anti-mouse Siglec-F (clone E50-2440) | BD Biosciences | Cat#: 552126 RRID: AB_394341 |
| PE anti-mouse F4/80 Antibody (clone BM8) | BioLegend | Cat#: 123110 RRID: AB_893486 |
| APC/Fire™ 750 anti-mouse CD19 Antibody (clone 6D5) | BioLegend | Cat#: 115558 RRID: AB_2572120 |
| APC anti-mouse IL-6 Antibody (clone MP5-20F3) | BioLegend | Cat#: 504508 RRID: AB_10694868 |
| Brilliant Violet 605™ anti-mouse TNF-α Antibody (clone MP6-XT22) | BioLegend | Cat#: 506329 RRID: AB_11123912 |
| PE/Cyanine7 anti-mouse CD64 (FcγRI) Antibody | BioLegend | Cat#: 139134 RRID: AB_2563904 |
| Brilliant Violet 711™ anti-mouse CD11c Antibody (clone N418) | BioLegend | Cat#: 117349 RRID: AB_2563905 |
| Brilliant Violet 510™ anti-mouse/human CD11b Antibody (clone M1/70) | BioLegend | Cat#: 101263 RRID: AB_2629529 |
| FITC anti-mouse CD170 (Siglec-F) Antibody (clone S17007L) | BioLegend | Cat#: 155504 RRID: AB_2750233 |
| PerCP/Cyanine5.5 anti-mouse anti-STAT1 Phospho (Ser727) Antibody (clone A15158B) | BioLegend | Cat#: 686416 RRID: AB_2734534 |
| Brilliant Violet 510™ anti-mouse F4/80 Antibody (clone BM8) | BioLegend | Cat#: 123135 RRID: AB_2562622 |
| Brilliant Violet 605™ anti-mouse CD11c Antibody (clone N418) | BioLegend | Cat#: 117334 RRID: AB_2562415 |
| APC anti-mouse CD284 (TLR4) Antibody (clone SA15-21) | BioLegend | Cat#: 145406 RRID: AB_2562503 |
| PE anti-mouse CD284 (TLR4) Antibody (clone SA15-21) | BioLegend | Cat#: 145404 RRID: AB_2561874 |
| APC/Cyanine7 anti-mouse TNF-α Antibody (clone MP6-XT22) | BioLegend | Cat#: 506344 RRID: AB_2565953 |
| PE-Cyanine7 anti-mouse c-MAF Monoclonal Antibody (clone sym0F1) | Invitrogen | Cat#: 25-9855-82 RRID: AB_2811795 |
| PE/Cyanine7 anti-mouse IFNAR-1 Antibody (clone MAR1-583) | BioLegend | Cat#: 127325 RRID: AB_2810385 |
| FITC CD282 (TLR2) Monoclonal Antibody (clone 6C2) | Invitrogen | Cat#: 11-9021-82 RRID: AB_465440 |
| TruStain FcX™ (anti-mouse CD16/32) Antibody (clone 93) | BioLegend | Cat# 101320 RRID: AB_1574975 |
| Ultra-LEAF™ Purified anti-mouse CD210 (IL-10 R) Antibody | BioLegend | Cat# 112710 RRID: AB_11149684 |
| FITC anti-mouse CD14 Antibody (clone Sa14-2) | BioLegend | Cat#: 123307 RRID: AB_940578 |
| APC anti-mouse CD14 Antibody (clone Sa14-2) | BioLegend | Cat# 123311 RRID: AB_940574 |
| APC/Cyanine7 anti-mouse CD3 Antibody (clone 17A2) | BioLegend | Cat# 100222 RRID: AB_2242784 |
| Purified Rat IgG1, κ Isotype Ctrl Antibody (clone RTK2071) | BioLegend | Cat# 400402 RRID: AB_326508 |
| Brilliant Violet 785™ anti-mouse/human CD11b Antibody (clone M1/70) | BioLegend | Cat# 101243 RRID: AB_2561373 |
| Chemicals, Peptides, and Recombinant Proteins | | |
| Brefeldin A Solution (1,000X) | BioLegend | Cat# 420601 |
| Lipopolysaccharide (LPS)-EB Biotin | InvivoGen | Cat. code tlrl-lpsbiot |
| Streptavidin Fluoresbrite® YG Microspheres, 1.0µm | Polysciences | Cat# 24161-1 |
| Cyto-Fast™ Fix/Perm Buffer Set | BioLegend | Cat# 426803 |
| Paraformaldehyde Aqueous Solution (20%) EM Grade | Electron Microscopy Sciences | Cat# 15713-S |
| CD11b MicroBeads, human and mouse | Miltenyi Biotec | Cat# 130-049-601 |
| Pam3CSK4 Biotin | InvivoGen | Cat. code tlrl-bpms |
| Phosphate Buffer Saline (PBS) | Gibco | Cat# 10010-049 |
| RPMI 1640 Media | Millipore Sigma | Cat# 11875-119 |
| L-Glutamine (200 mM) | Gibco | Cat# 25030-081 |
| Penicillin-Streptomycin (5,000 U/mL) | Gibco | Cat# 15140-122 |
| Fetal Bovine Serum (FBS) | Biowest | Cat# S1620 |
| True-Phos™ Perm Buffer | BioLegend | Cat# 425401 |
| Recombinant Human M-CSF | Peprotech | Cat# 300-25 |
| Ethylenediaminetetraacetic Acid (EDTA) (0.5M Solution/pH 8.0) | Fisher Scientific | Cat# BP24821 |
| Zombie Violet™ Fixable Viability Kit | BioLegend | Cat# 423113 |
| Zombie UV™ Fixable Viability Kit | BioLegend | Cat# 423107 |
| Lipopolysaccharide (LPS) source species S. minnesota R595 | List Labs | Cat# 434 |
| 2'3'-cGAMP | InvivoGen | Cat. code tlrl-nacga23-5 |
| Pam3CSK4 | InvivoGen | Cat. code tlrl-pms |
| Sodium Azide | Fisher Scientific | Cat# S227I-25 |
| Tris Base | Fisher Scientific | Cat# BP152-500 |
| LB Broth | Fisher Scientific | Cat# 244620 |
| LB Agar Kanamycin-50 Plates | Sigma-Aldrich | Cat# L0543 |
| DMEM (Dulbecco's Modified Eagle Medium) | Gibco | Cat# 11965-092 |
| HEPES (1 M) | Gibco | Cat# 15630-080 |
| Sodium Pyruvate (100 mM) | Gibco | Cat# 11360-070 |
| MEM Amino Acids Solution (50X) | Gibco | Cat# 11130-051 |
| Trypsin-EDTA (0.25%), phenol red | Gibco | Cat# 25200-056 |
| Poly-L-Lysine Solution (0.01%) | Sigma-Aldrich | Cat# A-005-C |
| TRIzol™ Reagent | Invitrogen | Cat# 15596018 |
| Chloroform | Thermo Scientific Chemicals | Cat# J67241.K2 |
| Isopropanol | Fisher Scientific | Cat# AC327270010 |
| PEI MAX® - Transfection Grade Linear Polyethylenimine Hydrochloride (MW 40,000) | Polysciences | Cat# 24765-100 |
| Polybrene Infection / Transfection Reagent | Millipore Sigma | Cat# TR-1003-G |
| Puromycin | InvivoGen | Cat. code ant-pr1 |
| Recombinant Mouse IFN-β1 (carrier-free) | BioLegend | Cat# 581306 |
| Recombinant Murine IL-10 | Peprotech | Cat# 210-10 |
| TaqMan™ Fast Universal PCR Master Mix (2X), no AmpErase™ UNG | Applied Biosystems | Cat# 4352046 |
| eBioscience™ Foxp3 / Transcription Factor Staining Buffer Set | Invitrogen | Cat# 00-5523-00 |
| Bovine Serum Albumin Solution (7.5% in DPBS) | Sigma-Aldrich | Cat# A8412 |
| Resiquimod (R848) | Sigma-Aldrich | Cat# SML0196 |
| GlycoBlue™ Coprecipitant (15 mg/mL) | Invitrogen | Cat# AM9515 |
| Kanamycin solution from *Streptomyces kanamyceticus* | Sigma Aldrich | Cat# K0254 |
| Critical Commercial Assays | | |
| Mouse TNF-alpha DuoSet ELISA | R&D Systems | Cat# DY410 |
| Mouse IL-6 DuoSet ELISA | R&D Systems | Cat# DY406 |
| Mouse IL-10 DuoSet ELISA | R&D Systems | Cat# DY417 |
| GeneJET Plasmid Miniprep Kit | Thermo Scientific | Cat# K0502 |
| QIAGEN Plasmid Maxi Kit (10) | QIAGEN | Cat# 12162 |
| Zombie Violet™ Fixable Viability Kit | BioLegend | Cat# 423113 |
| Zombie UV™ Fixable Viability Kit | BioLegend | Cat# 423107 |
| RNA to cDNA EcoDry™ Premix (Oligo dT) | TaKaRa | Cat# 639543 |
| RNA 6000 Pico Kit | Aligent | Cat# 5067-1513 |
| SMART-Seq v4 Ultra Low Input RNA Kit | TaKaRa | Cat# 634889 |
| Experimental Models: Cell Lines | | |
| 293T | ATCC | CRL-3216 |
| ProClone™ Competent Cells | ABM | Cat# E003 |
| Experimental Models: Organisms/Strains | | |
| Mouse: C57BL/6J | The Jackson Laboratories | Cat# 000664 In house breeding |
| Oligonucleotides | | |
| *Ifnb1* (6-FAM probe) | Integrated DNA Technologies | Assay ID:  Mm.PT.58.30132453.g |
| *Ifit3* (Cy5 Probe) | Integrated DNA Technologies | Assay ID: Mm.PT.58.33537107 |
| *Gapdh* (HEX Probe) | Integrated DNA Technologies | Assay ID: Mm.PT.39a.1 |
| *Il10* (Cy5 Probe) | Integrated DNA Technologies | Assay ID: Mm.PT.58.13531087 |
| *Maf* (FAM Probe) | Integrated DNA Technologies | Assay ID: Mm.PT.58.42145949.g |
| Recombinant DNA | | |
| MAF Lentiviral Vector (Mouse) (CMV) (pLenti-GIII-CMV) | ABM | Cat# 27897064 |
| pLenti-III-Blank Vector | ABM | Cat# LV587 |
| pCMV-VSV-G | Addgene | Plasmid #8454 |
| psPAX2 | Addgene | Plasmid #12260 |
| Software and Algorithms | | |
| FlowJo version 10 | BD Biosciences | N/A |
| R version 3.16 | R Development Core Team | <http://www.r-project.org/> |
| Adobe Illustrator | Adobe.com | N/A |
| Prism 10 | GraphPad | N/A |
| Python version 3.11.7 | Python Software Foundation | [http://www.python.org](http://www.python.org/) |
| Other | | |
| BD DUAL LSRFortessa (5 laser) | BD Biosciences | N/A |
| CFX Opus 96 Real-Time PCR System | Bio Rad | #12011319 |
| LC Sprint^®^ Reusable Nebulizer | PARI | Part No. 023F35 |
